# Hangover regulates gene expression by limiting NSL-mediated H4K16 acetylation

**DOI:** 10.1101/2025.04.22.649938

**Authors:** Jonathan Lenz, Laura Schmelzer, Ignasi Forné, Andrea Nist, Axel Imhof, Thorsten Stiewe, Alexander Brehm

## Abstract

The RNA-binding protein hangover is essential for several stress responses in *Drosophila melanogaster*. Here, we discover a novel function of hangover in the regulation of gene expression. Hangover binds to more than 2.000 genes in the *Drosophila* genome and modulates transcription. We identify a diverse set of chromatin regulators as hangover interactors, including NSL, dMec, Sin3A, dREAM and Ino80. Among these, the non-specific lethal complex (NSL) is the most prominent one. We show that hangover attenuates NSL-mediated H4K16 acetylation at transcriptional start sites to downregulate gene expression.

Our work uncovers novel roles for hangover in epigenetic gene regulation and suggests that it coordinates the function of multiple chromatin regulators.

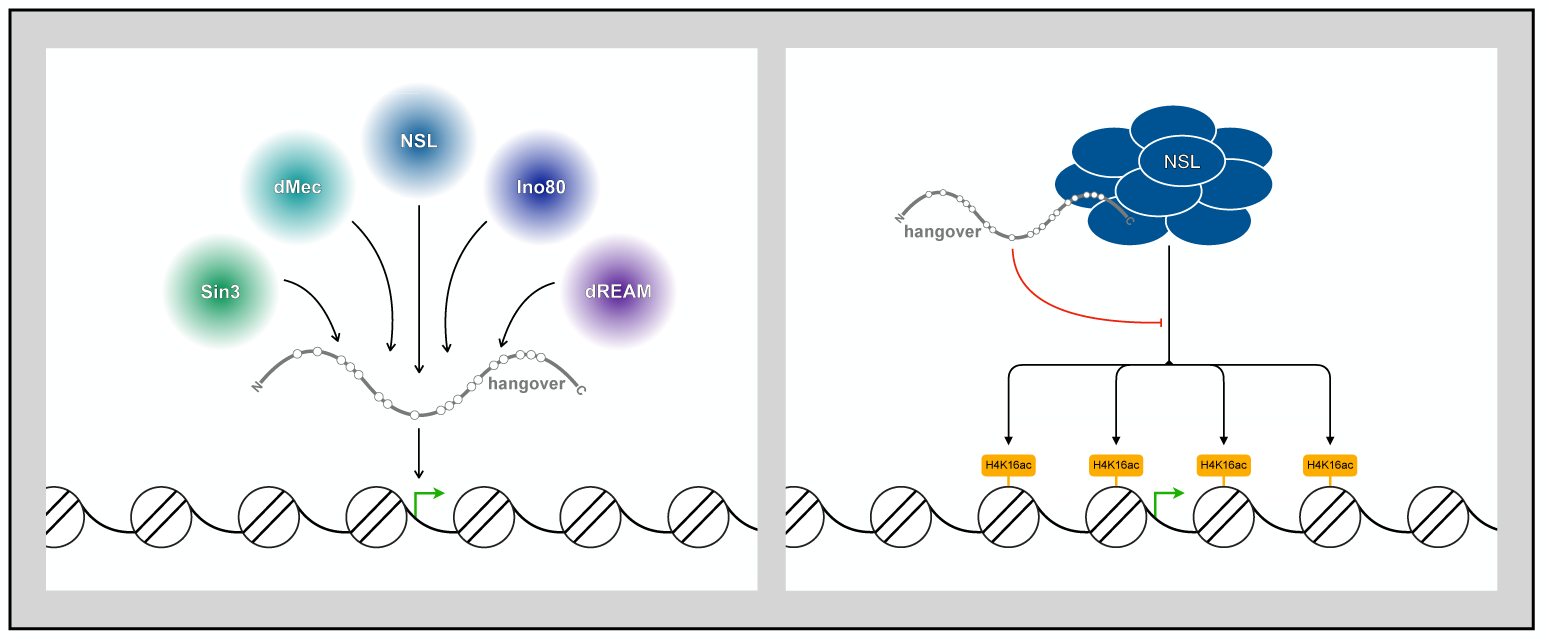

## Introduction

Cells have to respond to a variety of stimuli. These include physiological stimuli mediated by signaling molecules or cell-to-cell contacts as well as chemical and physical stresses. An adequate response includes modification of their gene expression profile and is often executed at multiple levels. Well studied cases of this are the heat shock response as well as the response to changes in nutrient availability via the mammalian target of rapamycin (mTOR) pathway. These signaling cascades not only result in adaptations to the cells transcriptome and chromatin landscape, but also adjust its mRNA translation efficiency (Teves et al., 2011; Guertin et al., 2010; Teves et al., 2013; Panniers et al., 1994; Elvira et al, 2020; Saxton et al., 2017).

The *Drosophila melanogaster* multi zinc finger protein hangover (hang) regulates responses to oxidative stress, heat shock and ethanol exposure. Fruit flies grow tolerant towards continual alcohol exposure in order to maintain postural control in high ethanol environments. As reflected in its name, hangover mutants fail to develop tolerance after repeated ethanol exposure resulting in impaired movement (Scholz et al., 2005). Hangover is expressed in the nuclei of motoneurons and directs the development of the neuromuscular junction. It regulates transcript levels in adult fly heads to enable an adequate stimulus response (Schwenkert et al., 2008, Ruppert et al., 2017). Its role in these pathways has primarily been studied using behavioral readouts and survival analyses of loss of function animals. However, little is known about its molecular activity.

The hangover protein contains no less than 18 C_2_H_2_-type zinc fingers. Using two matrin-type zinc fingers it binds and stabilizes the mRNA of the cAMP phosphodiesterase dunce (Ruppert et al., 2017). This suggests that one mechanism by which hangover regulates transcript levels is through RNA stabilization. C_2_H_2_ zinc fingers are prevalent domains among chromatin-regulating proteins, particularly transcription factors, where they typically mediate associations with DNA and transcriptional effectors (Cassandri et al., 2017; Brayer et al., 2008). Taken together with its nuclear localization and its ability to regulate mRNA levels it is conceivable that, in addition to its RNA stabilization function, hangover might impact on transcription.

Here, we identify the hangover interactome in *Drosophila* S2 cells by combining genomic epitope-tagging via CRISPR/Cas9 and a systematic proteomics approach. We uncover a large number of chromatin complexes that associate with hangover. ChIP sequencing reveals that hangover binds to chromatin and occupies 5’ regions of actively transcribed genes. We use RNA-seq to show that hangover predominantly represses mRNA expression in S2 cells. These observations position hangover as a transcriptional regulator.

To shed light on the mechanism by which hangover influences transcription, we further explore the interplay with its major interactor, the non-specific lethal (NSL) complex. We compared hangover-and NSL-dependent transcriptomes and find an antagonistic influence of both on gene expression. We investigate the NSL-catalyzed histone mark, acetylated lysine 16 in histone 4 (H4K16ac), in hangover knock-out cells using ChIP sequencing. This identified transcriptional start sites that exhibit increased H4K16ac in absence of hangover. These are associated with elevated expression and are linked to neuronal function and stimulus responses. We propose that hangover regulates the mRNA output of genes by modulating H4K16ac levels.

Our findings show that hangover not only regulates mRNA stability, but also impacts on chromatin modification as well as transcription in cooperation with versatile chromatin regulators.

## Results

### Hangover interacts with chromatin-regulating proteins

We identified hangover-interacting proteins in order to shed light on its molecular function. To this end, we used a CRISPR/Cas9-based approach to introduce GFP-or FLAG-tag sequences at the 3’ end of the *hangover* gene in S2 cells. This ensures the expression of a C-terminally tagged protein from its endogenous locus. Insertion of the epitope tag sequences was verified by PCR on genomic DNA and Western blot (**Suppl. Fig. 1A & B**). Subcellular fractionation showed that hangover is a nuclear protein in S2 cells. (**Suppl. Fig. 1C**). We performed ⍺-GFP and ⍺-FLAG immunoprecipitation on nuclear extracts from control cells and cells expressing hang-GFP or hang-FLAG. SDS-PAGE and silver staining of the precipitated material revealed the coprecipitation of several proteins (**Figure 1A**). Upon size exclusion chromatography (SEC) we detected hangover in fractions of high molecular weight, suggesting that it is part of larger protein complexes (**Suppl. Fig. 1D**). The hangover elution behaviour was not altered upon Benzonase digestion which indicates that interactions with nucleic acids are not critical for hangover to be incorporated into high molecular weight complexes.

**Figure 1.**
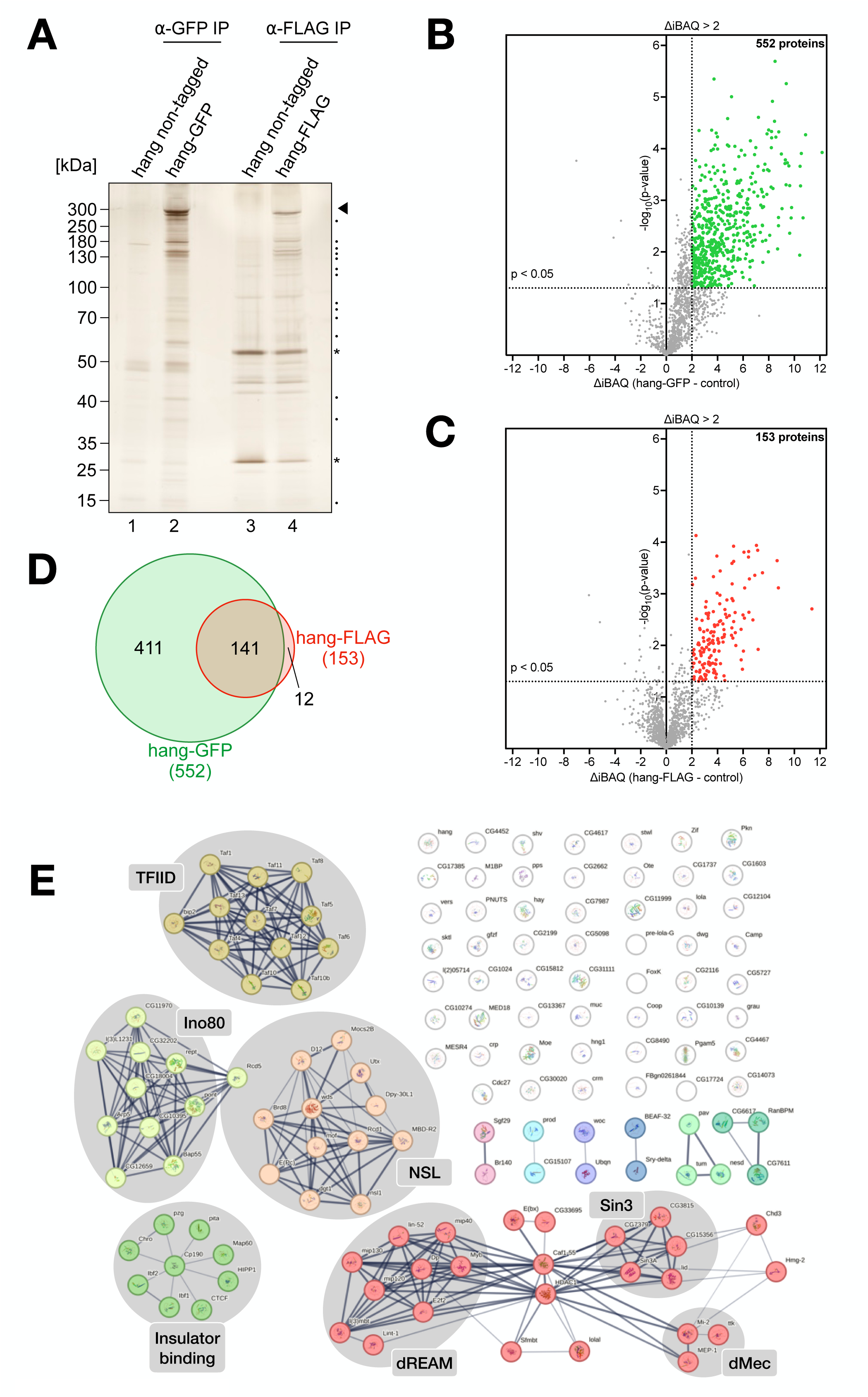
A. Silver-stained polyacrylamide gel of ⍺-GFP (lane 1 & 2) and ⍺-FLAG (lane 3 & 4) immunoprecipitates (IPs) from nuclear extracts of control S2 cells (hang non-tagged lanes 1 & 3) and cells expressing hang-GFP (lane 2) and hang-FLAG (lane 4). Molecular weight marker bands are indicated on the left. Arrowhead points to hangover, dots to polypeptides that are specifically enriched in hang-GFP and hang-FLAG IPs, respectively. Asterisks indicate heavy and light chain of the FLAG antibody. **B & C** IP-MS/MS results of ⍺-GFP (**B**) and ⍺-FLAG IP (**C**) represented as volcano plots. Proteins with a Δ log2 iBAQ enrichment > 2 and p<0.05 are marked in green (**B**) and red (**C**), respectively. The number of these proteins is indicated above. **D** Venn diagram showing the overlap of hang-GFP and hang-FLAG interacting proteins. **E** STRING interaction network of hangover high-confidence interactors after clustering. Lines between proteins indicate physical interactions. Line density represents the interaction confidence. Prominent chromatin-associated complexes are marked in grey.

We identified hangover-interacting proteins by analyzing hang-GFP and hang-FLAG coprecipitates with mass spectrometry (LC-MS/MS). Compared to control IPs from extracts containing non-tagged hangover, 552 proteins were enriched in hang-GFP precipitates and 153 proteins were enriched in hang-FLAG precipitates (**Figure 1B & C**). 141 proteins were enriched in both conditions which we refer to as “high-confidence interactors” (**Figure 1D**, Suppl. Table 2). These results support the notion that hangover does not operate as a monomer but cooperates with other proteins. In order to assign biological functions to hangover interactors we employed Gene Ontology (GO) analysis. The most significantly enriched GO terms were associated with chromatin and transcription (“chromatin organization”, “ATP-dependent chromatin remodeling”, “negative regulation of transcription”, “positive regulation of transcription”) (**Suppl. Fig. 1E**). In fact, 83 out of the 141 high-confidence interactors (58.9%) were assigned to one or more transcription-related GO term. This is unexpected since hangover has so far not been implicated in the regulation of transcription or chromatin. Using STRING interaction analysis, we sorted hangover high-confidence interactors into their respective protein complexes (**Figure 1E**). This identified several multi-subunit chromatin-regulators: Hangover interacts with chromatin remodelling complexes (Inositol-requiring mutant 80 (Ino80) and *Drosophila* MEP-1 complex (dMec)) and histone-modifying complexes (NSL and Swi-independent 3 (Sin3)). It contacts activators (TFIID) as well as repressors (*Drosophila* Rbf, E2F and Myb-interacting proteins (dREAM)) of transcription. Moreover, it interacts with a cluster of insulator-binding proteins.

We conclude that hangover associates with several chromatin-regulating protein complexes. Given the predominance of transcriptional regulators among its high-confidence interactors we hypothesize that hangover itself binds and regulates chromatin.

### Hangover binds to actively transcribed genes

In order to determine whether hangover associates with chromatin we performed chromatin immunoprecipitation (ChIP) of hangover in hang-GFP expressing cells. We analyzed two replicates by high throughput sequencing and identified more than 3000 binding sites in the genome of S2 cells that were shared between both replicates (**Figure 2A**). We determined the genomic distribution of hangover binding sites. Hangover signal was most strongly enriched at the 5’ end of genes with its highest binding intensity around 150 base pairs downstream of transcriptional start sites (**Figure 2B & C**). To determine if hangover associates with actively transcribed and/or repressed genes, we investigated the chromatin environment at hangover-bound regions. Hangover colocalizes with histone modifications associated with active transcription (H3K4me3 and H3K27ac). By contrast, its binding sites are mostly devoid of H3K27me3, a mark of polycomb-mediated gene repression. Hangover-bound regions also show high ATAC-seq signal indicating increased chromatin accessibility (**Figure 2D**). To quantify their transcriptional status, expression levels of hangover-occupied genes were determined. Genes underlying hangover binding sites exhibited significantly higher mRNA levels in comparison to all genes expressed in S2 cells (**Figure 2E**). Collectively, these observations demonstrate that hangover binds to actively transcribed genes.

**Figure 2.**
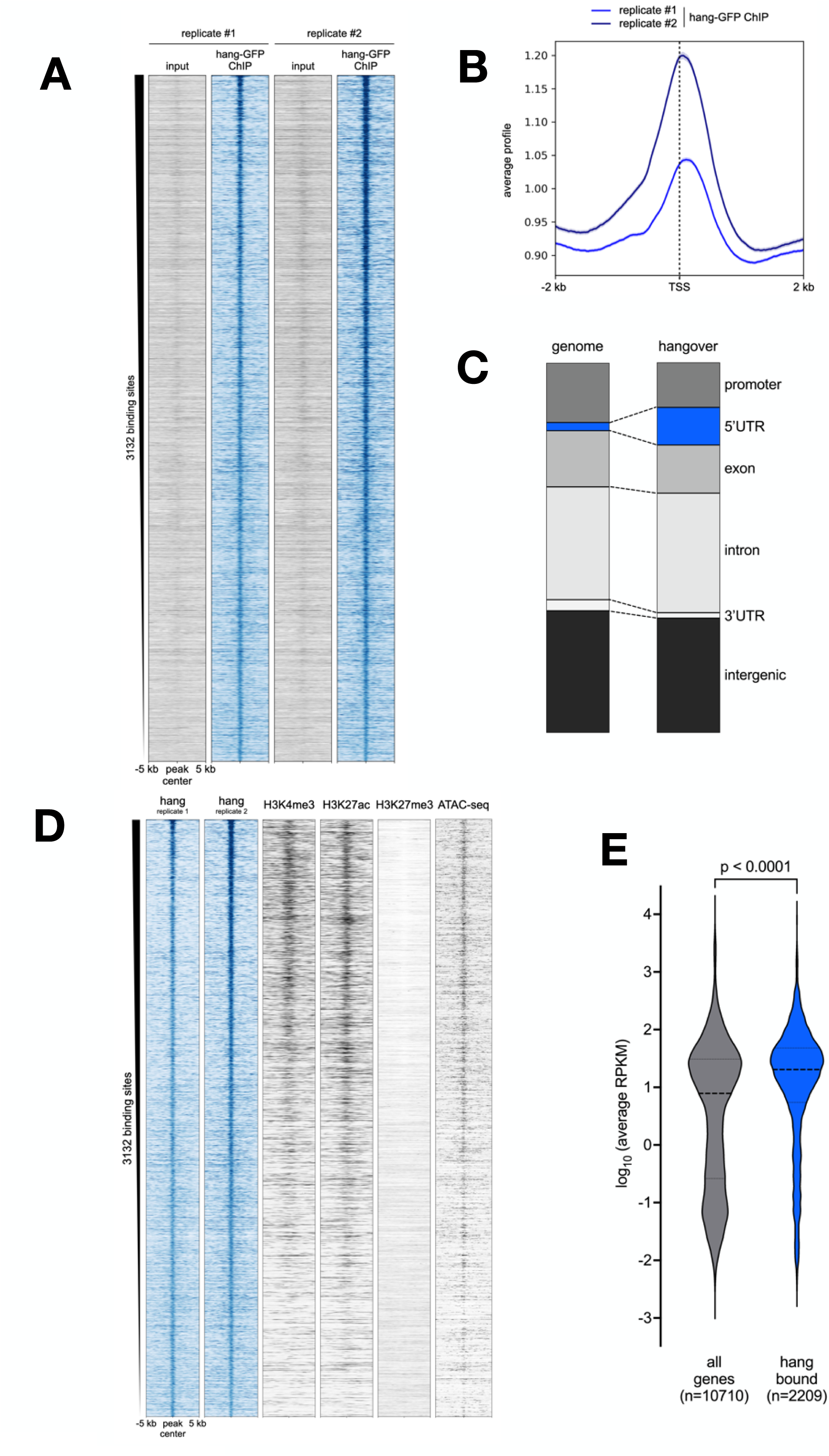
A. Heatmap of hang-GFP and input ChIP-seq signals within a 10 kb interval of hang-GFP binding sites. Results of two hang-GFP ChIP-seq replicates in S2 cells are shown. **B** Metagene analysis of hang-GFP ChIP-seq signal at TSS. Normalized hangover signal was evaluated within a 4 kb interval surrounding all genomic TSS. The two graphs represent two hang-GFP ChIP-seq replicates (light blue: replicate #1, dark blue: replicate #2). **C** Distribution of hang-GFP binding sites over genomic regions compared to the distribution of these regions in the genome. The most highly enriched region (5’ UTR) is indicated in blue. **D** Heatmap of hang-GFP ChIP-seq, histone modification ChIP-seq and ATAC-seq signals within a 10 kb interval of hangover binding sites. **E** Average transcript levels (average RPKM) of all genes expressed in S2 cells (grey) and genes bound by hangover (blue). Their distribution is shown as violin plots. Dashed lines represent the median and quartiles. Distributions were compared using two-tailed Mann-Whitney statistics.

### Hangover represses transcription

Its association with actively transcribed genes and its interaction with transcriptional regulators strongly suggest that hangover impacts on gene expression. In order to identify hangover-dependent transcripts we performed RNA sequencing after depletion of hangover from S2 cells using RNAi. We used two different double stranded RNA (dsRNA) sequences and confirmed effective depletion by Western blot (**Suppl. Fig. 3A**). RNA from cells transfected with control dsRNA (dsEGFP) and from cells transfected with dsRNA targeting hangover (dshang#1 and dshang#2) was analyzed by high throughput sequencing. Depletion of hangover with the first construct resulted in the upregulation of 356 genes, whereas 149 showed lower expression (**Figure 3A**). Depletion with the second construct led to significant derepression of 442 genes and decreased the levels of 243 transcripts (**Figure 3B**). Both data sets showed a high degree of correlation of the significantly differentially expressed genes (**Figure 3C**, R^2^=0.9414). 260 genes responded with an increased expression upon hangover depletion, whereas 65 genes were downregulated in both experiments (**Figure 3C**, Suppl. Table 3). Due to the higher number of genes that are repressed by hangover we conclude that hangover primarily represses transcription.

**Figure 3.**
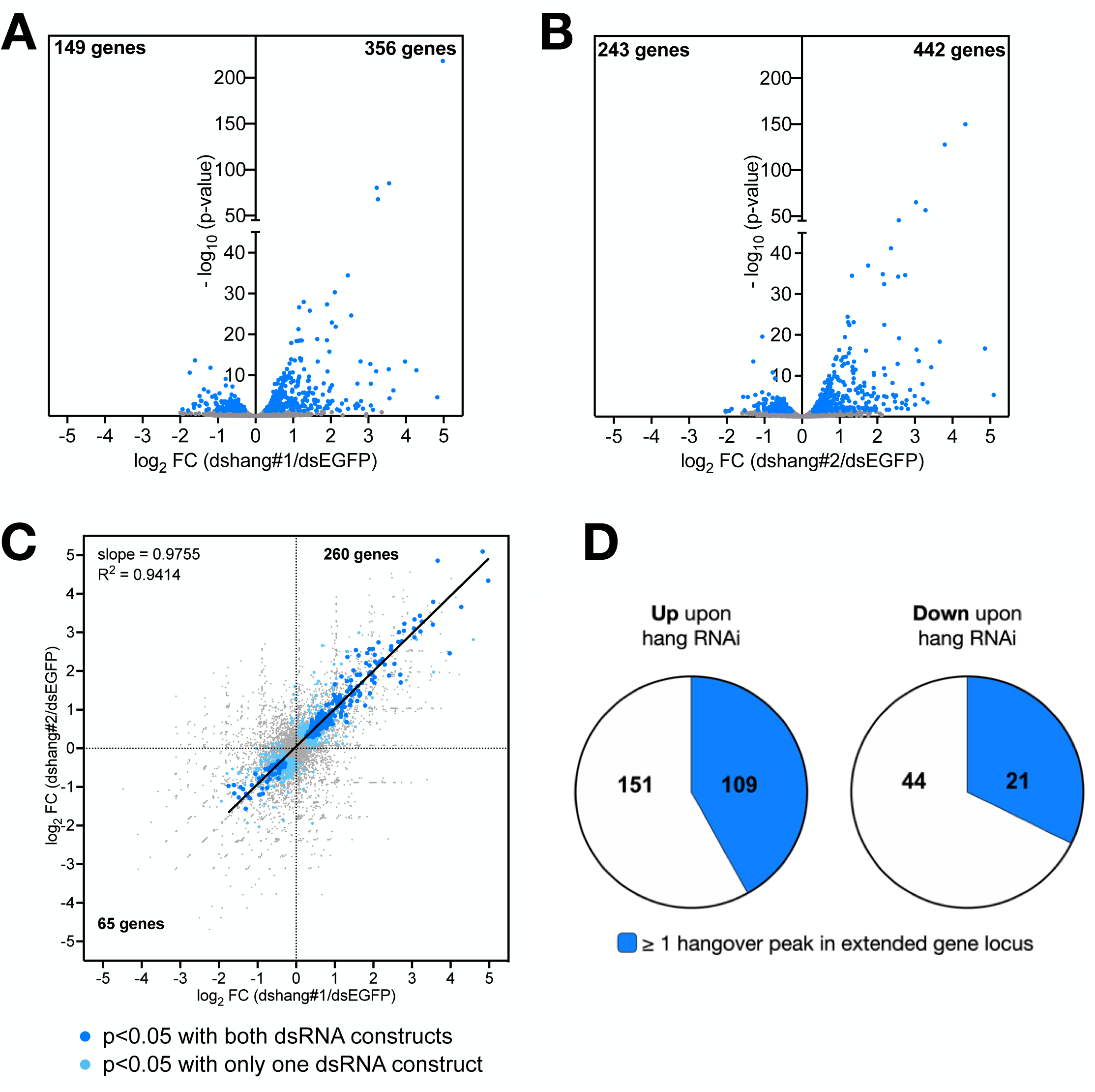
A &. **B** Volcano plots of genes deregulated upon RNAi-mediated depletion of hangover. Log_2_ fold changes of hangover dsRNA treated vs. dsEGFP treated cells are shown for the first (**A**, dshang#1) and the second RNAi construct (**B**, dshang#2). Genes with p<0.05 are marked in blue and their numbers are indicated above. **C** Correlation analysis of fold gene expression changes upon depletion with the two dsRNA constructs used to target hangover. Colors represent genes whose fold change (FC) is associated with a p-value < 0.05 in both (blue), only one (light blue) or neither condition (grey). Numbers of genes with p<0.05 in both conditions are indicated in the first and third quadrant. Linear regression of genes with p<0.05 in both conditions is indicated in the upper left. **D** Pie chart depicting hangover binding to hangover-repressed and hangover-dependent genes. Charts are separated by genes whose expression is repressed by (left) or dependent on (right) hangover. Blue sections represent genes that contain one or more hangover binding sites.

We examined hangover occupancy at hangover-dependent genes. 41.9% of genes (109 out of 260) that showed increased expression levels and 32.3% of genes (21 out of 65) that showed lower expression levels upon hangover depletion also contained a hangover binding site (**Figure 3D**). This suggests that hangover regulates transcription, at least in part, by directly contacting their gene loci. We propose that hangover employs the activity of its interacting chromatin regulators to modulate transcription.

### Hangover and NSL have opposite effects on gene expression

Hangover binds to chromatin, it interacts with transcriptional regulators and impacts on gene expression. Based on these observations we hypothesized that hangover contributes to shaping the chromatin environment at actively transcribed genes. To elucidate how hangover modulates transcription we focussed on the non-specific lethal (NSL) complex since NSL subunits were among the most highly enriched proteins in our hangover interactome analysis (**Suppl. Fig. 4A & B**, Suppl. Table 2). We confirmed the association of hangover with NSL by coimmunoprecipitation followed by Western blot. Presence of the NSL subunits nsl1, MBD-R2, Rcd5 and mof was observed in both hang-GFP and hang-FLAG precipitates (**Figure 4A**). Additionally, the bulk of the hangover protein coeluted with NSL complex components in size exclusion chromatography (**Suppl. Fig. 4C**). These findings confirm that NSL is the most prominent hangover interactor in S2 cells.

**Figure 4.**
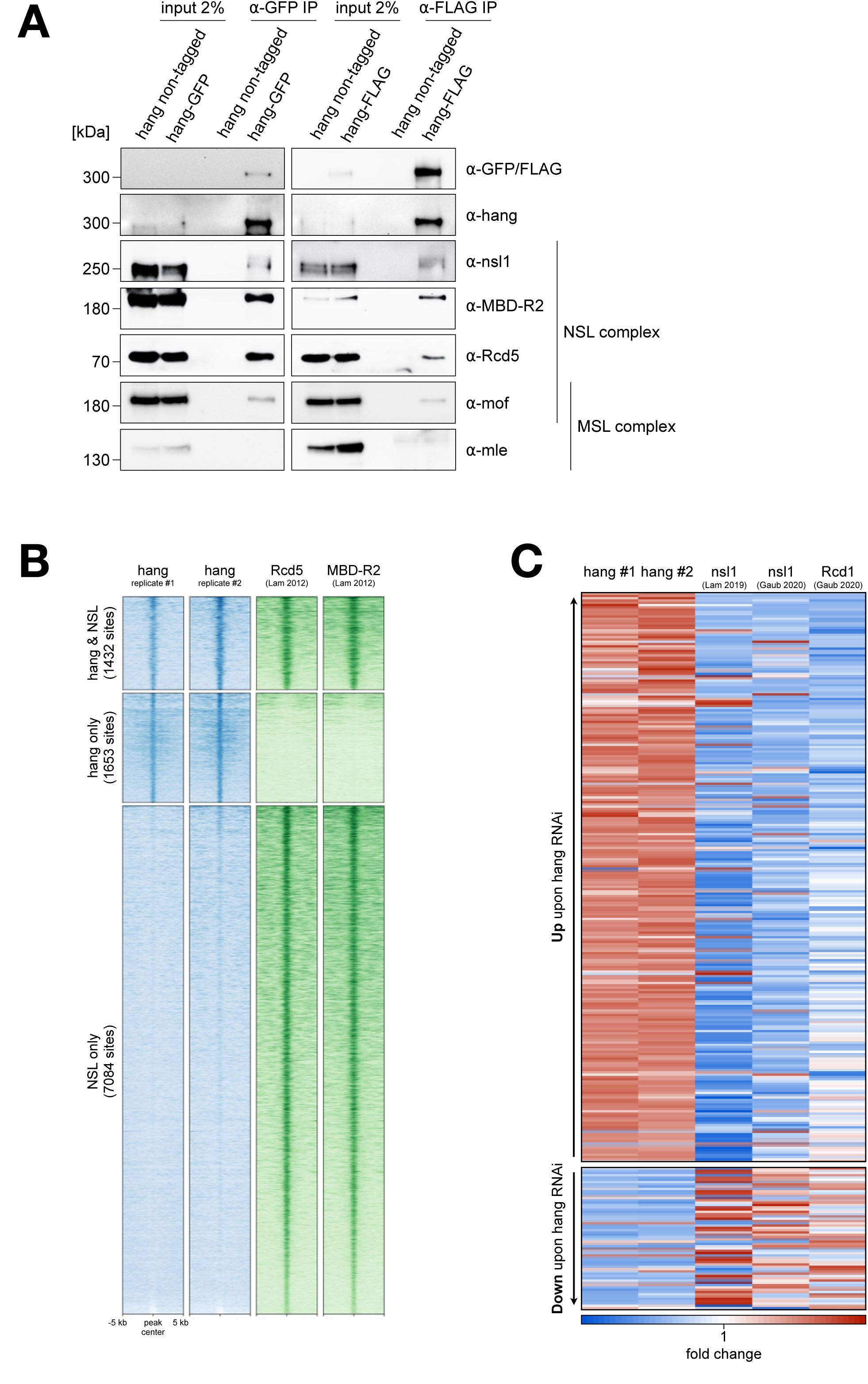
A. Coimmunoprecipitation of NSL complex subunits with hang-GFP (left panels) and hang-FLAG (right panels). ⍺-GFP and ⍺-FLAG IP was performed from nuclear extracts of control cells (hang non-tagged) and cells expressing hang-GFP or hang-FLAG, respectively. Immunoprecipitates were verified using GFP, FLAG and hangover antibodies and probed with antibodies against NSL and MSL subunits as indicated on the right. Molecular weight markers are indicated on the left. **B** Heatmap of ChIP-seq signals at hangover and NSL binding sites. Signal of both hangover-GFP ChIP-seq replicates (blue) and the NSL complex subunits Rcd1 and MBD-R2 (green) are plotted in a 10 kb window surrounding peak centers. Heatmap is divided according to co-occupied (top), hangover only (middle) and NSL only (bottom) regions. **C** Heatmap depicting fold changes of hangover-regulated genes upon depletion of hangover (lanes 1 & 2) and the NSL complex subunits nsl1 and Rcd1 (lanes 3-5). Top panel depicts 260 genes with increased expression, bottom panel depicts 65 genes with decreased expression upon hangover depletion. Values are scaled by row and sorted according to FC upon hangover RNAi.

NSL is a ubiquitous chromatin regulator that contains the histone acetyltransferase mof. By acetylating histone 4 at lysine 16 (H4K16) it activates gene transcription (Sheikh et al., 2019). The catalytic subunit of NSL, mof, is also found in a second protein assembly, the male-specific lethal (MSL) complex, which is involved in dosage compensation (Gelbart et al., 2009). To test whether hangover might also interact with MSL we probed hang-GFP and hang-FLAG precipitates for the occurrence of the MSL subunit mle. We did not detect mle as an interactor of hangover (**Figure 4A**). Additionally, the MSL subunits msl-1, msl-2 and msl-3 were not part of hangover high-confidence interactors (**Suppl. Fig 4A & B**). This indicates that hangover specifically contacts NSL and not MSL.

Given that hangover and NSL interact in nuclear extracts we asked whether they also colocalize on chromatin. We compared hangover binding sites to published ChIP sequencing data of the NSL subunits Rcd1 and MBD-R2 (Lam et al., 2012). 46.4% of hangover binding sites (1432 out of 3085) colocalized with both Rcd1 and MBD-R2, whereas 53.6% (1653 out of 3085) did not show substantial enrichment of NSL (**Figure 4B**). Moreover, there was a large number of additional NSL binding sites (7084) lacking significant hangover signal. This is not surprising since NSL is an abundant complex that regulates the expression of thousands of genes (**Suppl. Fig. 4D-F**). We conclude that NSL and hangover not only interact in the soluble nuclear fraction, but also share binding sites on chromatin.

Our RNA sequencing analysis suggests that hangover predominantly represses gene expression (**Figure 3**). However, the NSL complex is an activator of transcription. To investigate the apparently opposing effects on transcription we elucidated the influence of NSL on the expression of hangover-regulated genes. To this end we analyzed RNA sequencing data upon depletion of nsl1 and Rcd1 (**Suppl. Fig. 4D-F**; Lam et al., 2019; Gaub et al., 2020). Gene expression changes were investigated over the set of genes that responded to hangover depletion. Intriguingly, genes that were repressed by hangover required NSL for their activation. On the contrary, genes downregulated upon hangover depletion tended to respond with higher expression to the loss of NSL (**Figure 4C**). This opposite influence on genes was most strikingly observed at hangover-repressed genes upon depletion of nsl1 (**Suppl. Fig. 4G**). We conclude that hangover and NSL indeed have opposite effects on gene expression.

### Loss of hangover causes local changes in H4K16ac

We hypothesized that hangover might antagonize NSL function. To test this hypothesis, we used CRISPR/Cas9 to generate hangover knock-out cells (hang ko) by inserting a GFP expression cassette in the *hangover* locus of hang-FLAG expressing cells (hang wt). The GFP cassette was inserted into codon 2 of the hangover gene thereby disrupting its open reading frame. This setup allowed us to monitor remaining hangover expression by Western blot against the FLAG epitope. Insertion was confirmed by PCR on genomic DNA (**Suppl. Fig. 5A**). Hangover ko cells showed no remaining hang-FLAG signal in Western blot but were positive for GFP expression (**Suppl. Fig. 5B**) confirming absence of hangover in these cells. To explore the effects of hangover on mof-catalyzed histone acetylation we examined the genomic distribution of H4K16ac using ChIP sequencing in hangover wt and ko cells. We quantified the spike-in normalized genome-wide H4K16ac ChIP-seq signal (**Suppl Fig. 5C**). We did not observe global changes in H4K16ac upon hangover knock-out. This is not unexpected since, in male S2 cells, the majority of H4K16ac is deposited by the MSL complex and resides on the X chromosome (Gelbart et al., 2009). To investigate locus-specific effects we focused our analysis on the transcriptional start sites (TSS) of genes. We identified 192 TSS that gained H4K16 acetylation upon hangover knock-out and 237 TSS that showed decreased H4K16ac (**Figure 5A & B; Suppl. Fig. 5D**). These changes were especially pronounced on TSS that showed low H4K16ac levels in wild type cells. It is noteworthy that loss of hangover changes H4K16 acetylation on a narrow subset of TSS. This corresponds well to the limited number of genes that are deregulated upon depletion of hangover compared to the thousands of genes that are controlled by NSL (**Figure 3; Suppl. Fig. 4 D-F**). Importantly, TSS on the X chromosome were affected to a lower extent than TSS on other chromosomes further supporting the notion that altered H4K16ac levels in hangover knock-out cells are likely attributed to NSL rather than to MSL function (**Suppl. Fig. 5E**).

**Figure 5.**
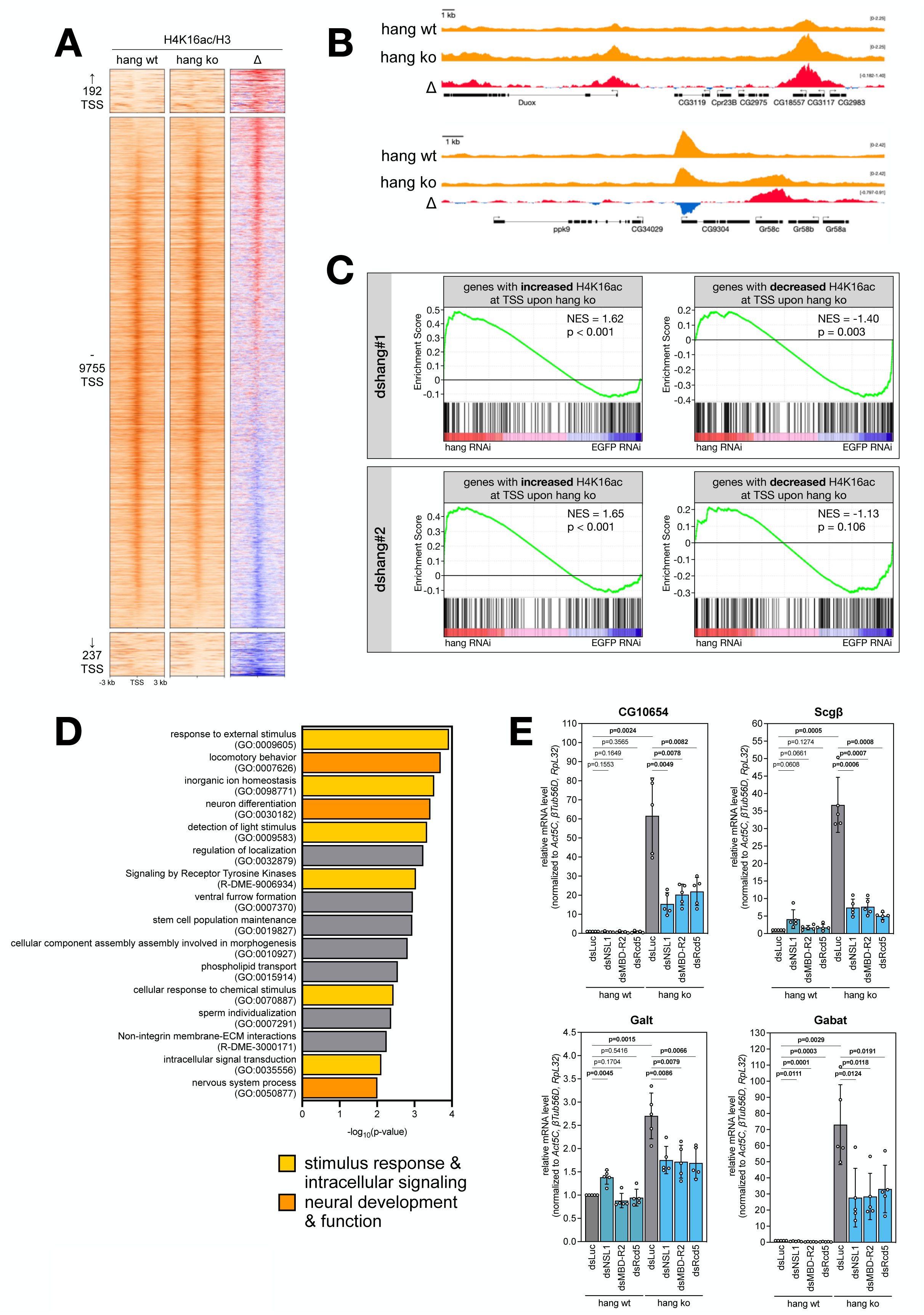
A. Heatmap depicting H4K16ac ChIP-seq signal in a 6 kb interval surrounding TSS in wt (left panel) and hangover ko cells (middle panel). Difference between wt and ko signal is represented in the right panels; red denotes increased and blue decreased H4K16ac ChIP signal. Most highly affected TSS are shown on top (192 TSS with increased H4K16ac in hangover ko) and bottom (237 TSS with increased H4K16ac in hangover ko). **B** Genome browser snapshots of exemplary regions containing differentially H4K16 acetylated TSS upon hangover ko. H4K16ac signal is shown in hangover wt (top) and ko cells (bottom), as well as difference between both cell lines (bottom, red: increase, blue: decrease). **C** Gene Set Enrichment Analysis of RNA-seq data from hangover depleted cells (top panel: RNAi with construct #1, bottom panel: RNAi with construct #2). Tested gene sets were comprised of genes with increased (left) and decreased (right) H4K16ac at their TSS in hangover ko cells. Normalized enrichment scores (NES) and p-values are indicated. **D** Gene Ontology term analysis of genes with decreased H4K16ac signal at their TSS in hangover ko cells. GO terms associated with stimulus response & intracellular signaling are highlighted in yellow. GO terms associated with neural development & function are highlighted in orange. **E** RTqPCR analysis of selected genes with increased H4K16ac at their TSS in hang ko cells. Expression was evaluated following RNAi in wt and hangover ko cells upon control transfection (dsLuc) or depletion of NSL complex subunits (dsNsl1, dsMBD-R2, dsRcd5). Values are expressed as FC over control transfection in wt cells and are normalized to expression of *Act5C*, *βTub56D* and *RpL32*. Error bars represent standard deviation from independent transfections (n=5), which are depicted as dots. Distributions were compared using unpaired, Welch corrected T statistics and p-values are indicated above.

In order to examine whether the acetylation change upon hangover loss is predictive of an expression change of the underlying gene, we used Gene Set Enrichment Analysis (GSEA) on RNA sequencing data upon hangover RNAi. We compiled sets of genes that exhibited higher or lower H4K16ac levels at their TSS in hangover knock-out cells. Genes with an increase in H4K16ac showed significantly higher expression levels upon hangover depletion (**Figure 5C**). This suggests that hangover-mediated repression is, in part, realized by restricting acetylation of H4K16. The expression levels of genes that exhibited decreased H4K16ac at their TSS were not significantly affected upon hangover RNAi, indicating that here lower H4K16ac does not cause detectable transcriptional changes.

We used Gene Ontology analysis to gain insight into the function of those genes that gained H4K16ac in hangover knock-out cells. Remarkably, we recovered GO terms associated with neural development and function (“locomotory behavior”, “neuron differentiation”, “nervous system process”) as well as gene groups linked to stimulus responses and intracellular signaling (“response to external stimulus”, “detection of light stimulus”, “Signaling by Receptor Tyrosine Kinases”, “cellular response to chemical stimulus”, “intracellular signal transduction”) (**Figure 5D**). This indicates that hangover particularly limits acetylation of gene promoters linked to response to environmental signals and neuronal function.

The observations above evoke a scenario in which hangover restricts the activity of NSL and this way prevents transcriptional activation. We chose four exemplary genes with increased H4K16ac in hangover knock-out cells to test this model: *CG10654*, *Scgβ*, *Galt* and *Gabat* (**Suppl. Fig. 5D**). Three of these genes are highly expressed in adult brains (FlyAtlas2; Leader et al., 2018; Krause et al., 2022) and the central nervous system of larvae (modENCODE): The zinc finger transcription factor CG10654, the sarcoglycan complex component Scgβ and the GABA transaminase Gabat. Using RNAi we depleted the NSL complex subunits nsl1, MBD-R2 and Rcd5 in hangover wt and ko cells followed by RTqPCR (**Figure 5E; Suppl. Fig. 5F**). All four genes are induced upon loss of hangover. In wild type cells depletion of NSL components had little or no effect on their expression. In hangover ko cells, however, the increased mRNA levels caused by hangover loss were significantly impaired upon NSL RNAi, resulting in decreased expression (**Figure 5E**). This demonstrates that the activation of transcription upon hangover depletion depends on the NSL complex.

We conclude that hangover represses transcription by limiting NSL-mediated H4K16 acetylation at a number of TSS associated with neural development and stimulus response.

## Discussion

### Hangover is a transcriptional regulator

We have shown that the multi zinc finger protein hangover binds to chromatin and is predominantly localized at the 5’ end of genes. This region harbours regulatory sequences crucial for gene transcription such as promoters, the TSS as well as RNA polymerase II pausing sites. More than 300 genes are deregulated upon hangover depletion and a substantial number of their gene loci are occupied by hangover. We therefore propose that hangover regulates gene expression, in part, by directly influencing transcription. This is further supported by the fact that hangover knock-out results in local changes to H4K16ac levels, a histone mark linked to transcriptional activation.

We have shown that hangover occupies active genes, however, it predominantly represses transcription. Moreover, it binds to both activating and repressive chromatin-regulating proteins. Therefore, we consider it likely that hangover is required to modulate the mRNA output of active genes rather than establishing silent heterochromatin.

Hangover directly binds and stabilizes mRNA in neuronal tissue (Ruppert et al., 2017). The novel chromatin functions of hangover identified here represent a second tier of regulating mRNA levels. Thus, hangover adds to the growing number of transcription factors that are capable of binding to both chromatin and RNA (Oksuz et al., 2023).

### Hangover binds to multiple chromatin regulators

The majority of hangover interactors are regulators of chromatin and transcription. Our list of interactors also includes the Polycomb group factor Sfmbt which has previously been shown to bind to hangover (Erokhin et al., 2021). Prominent hangover-associated complexes include dMec and Ino80 complexes which contain ATPases responsible for nucleosome remodelling (Kunert et al., 2009; Clapier et al., 2009). The Sin3 complex contains the histone deacetylase RPD3 and is linked to transcriptional repression (Spain et al., 2010). The NSL complex, on the other hand, activates gene expression by acetylating histones with its catalytic subunit mof (Sheikh et al., 2019). We recovered almost all subunits of the general transcription factor TFIID, suggesting a broader role for hangover in transcription initiation.

Its chromatin association and its interaction with effectors of gene expression evoke a scenario in which hangover recruits these factors to their sites of action. The simultaneous interaction with multiple regulators might allow hangover to direct combinations of their activities to certain gene loci. In doing so, hangover might serve as a hub or scaffold onto which chromatin-regulating complexes can assemble (Partridge et al., 2020; Malovannaya et al., 2011).

Additionally, hangover interacts with insulator-binding proteins (e.g. CTCF, Cp190 and Ibf1/2) indicating that it might partake in chromatin insulation or even spatial genome organization (Kahn et al., 2023; Bhattacharya et al., 2024).

### Hangover counteracts NSL mediated H4K16 acetylation

The most prominent binding partner of hangover in S2 cells is the NSL complex. We identified a group of genes that are repressed by hangover and activated by NSL. We therefore consider a recruitment model in which hangover tethers NSL to their shared binding sites unlikely. Rather, hangover antagonises NSL function. Here, we can envision two mechanisms: (1) hangover might counteract NSL binding to a subset of NSL-bound TSS; (2) hangover might modulate the catalytic activity of NSL.

The NSL complex is classically known to maintain expression of housekeeping genes (Lam et al., 2012). However, it has also been linked to stress-induced transcription (Sheikh et al., 2016). Since hangover is required for appropriate stress responses in adult flies we propose that this function partly relies on modulating NSL activity. This hypothesis is supported by the fact that genes with elevated H4K16ac levels at their TSS upon hangover knock-out are associated with stimulus response and intracellular signal transduction. Thus, it is plausible that phenotypic alterations in hangover loss of function animals (i.e. impaired stress response, neuronal malformations) might in part be caused by altered H4K16ac levels.

### Concluding remarks

Environmental stimuli and stresses demand a complex genomic response. Multi zinc finger proteins seem particularly suited to coordinate this task, since they have the potential to interact with RNA, chromatin and transcriptional regulators (Oksuz et al., 2023, Schmitges, Radovani, et al., 2016). The *Drosophila* transcription factor hangover is a striking example of this: it directly interacts with RNA (Ruppert et al., 2017), binds to thousands of sites on chromatin and contacts many cofactors with diverse chromatin-related activities. How these different binding activities are integrated in response to external stimuli is still an open question.

## Methods

### Cell culture

*Drosophila melanogaster* Schneider 2 cells with stable Cas9 expression (S2[Cas9] cells, clone 5-3) were kindly provided by Klaus Förstemann (Böttcher et al., 2014) and served as parental line for all cell lines generated in this study. Cells were cultivated in Schneider’s *Drosophila* Medium (2172001, Gibco) supplemented with 10% (v/v) fetal bovine serum (FBS; F7424, Sigma) and 1% (v/v) Penicillin-Streptomycin (15140122, Gibco). Cells were grown under standard conditions at 26 °C.

### Genomic tagging and knockout using CRISPR/Cas9

Genetic modification of S2 cells was performed as previously described (Böttcher et al., 2014; Lenz et al., 2021). Endogenous tagging was carried out by inserting GFP-or FLAG-tag sequences at the 3’ end of the *hangover* gene locus, resulting in the expression of a C-terminally tagged protein. To generate knock-out cell lines, a GFP sequence was inserted directly downstream of the start codon in order to disrupt subsequent transcription of the hangover coding sequence.

To that end, Cas9-stable S2 cells (10^6^/ml) were transfected with dsRNA against key enzymes of NHEJ (Lig4) and MMEJ (mus308) (1 µg/ml each) to favour repair by homologous recombination. After three days, cells (1.5 x10^6^/ml) were transfected with two PCR-generated dsDNA constructs: the first one containing the sgRNA sequence under control of a U6 snRNA promoter and the second one providing the tag and a resistance marker flanked by 60 bp homology. Transfection was carried out using FuGENE HD (E2311, Promega). sgRNA targeting sequences were AGCCTGACTCCAAGTCAG (tagging of hangover isoform A) and ACAAAATGTGCGACGCTG (hangover knock-out). Homologous recombination repair constructs were amplified from the following plasmids: pMH3 (GFP-tag & Blasticidin resistance marker, Addgene #52528), pMH4 (2xFLAG-tag & Blasticidin resistance marker, Addgene #52529), pSK23 (GFP sequence & Puromycin resistance marker, Addgene #72851).

Four days after transfection, cells were transferred into selection medium (10 µg/ml Blasticidin (A11139, Gibco) or 5 µg/ml Puromycin (540411, Merck)). Selection antibiotics were kept present until non-resistant control cells declined or at least for 14 days). Monoclonal cell lines were generated by serial dilution and the insertion was verified by PCR on genomic DNA. Primers for amplification of tagging constructs and genotyping can be found in table S1.

### Chromatin Immunoprecipitation (ChIP)

10^8^ S2[Cas9] cells were fixed with 1% (v/v) Formaldehyde at room temperature (RT) for 10 min. The reaction was stopped with 240 mM Glycin for 10 min at RT. Cells were washed with PBS and then lysed in 1 ml ChIP Lysis buffer (50 mM Tris/HCl pH 8.0, 10 mM EDTA, 1% (w/v) SDS, 1 mM DTT). After 10 min on ice, chromatin was fragmented using a Bioruptor UCD-200TM-EX (Diagenode) supplied with ice water for 30 min (30 sec sonication at high power followed by 30 sec cool down). Lysate was cleared by centrifugation at 21,100 g for 30 min at 4 °C and stored at-80 °C. Fragment size was evaluated on Agarose/TAE gels by decrosslinking 50 µl of lysate overnight at 65 °C including RNase A (400 ng/µl; A3832, Applichem) and Proteinase K (400 ng/µl; 7528.1, Roth). 140 µl of lysate was diluted 1:10 with ChIP IP buffer (16.7 mM Tris/HCl pH 8.0, 1.2 mM EDTA, 167 mM NaCl, 1.1% (w/v) Triton X-100, 0.01% (w/v) SDS, 1 mM DTT) and pre-cleared for 1 h using 40 µl of Protein A Sepharose (nProtein A Sepharose 4 Fast Flow, 17-5280, GE Healthcare) blocked with 2 mg/ml BSA and 2% (w/v) fish skin gelatin. For ChIP of histone modifications, 14 µl (1%) of HEK cell chromatin was added as spike-in control.

Antibodies were added to pre-cleared chromatin and incubated for 3 hours at 4 °C with rotation. Immune complexes were precipitated by adding 45 µl blocked Protein A Sepharose and incubation over night at 4 °C with rotation. GFP-tagged proteins were immunoprecipitated using 25 µl of blocked GFP-Trap Agarose (gta, Proteintech).

Protein-coupled resin was washed thrice with ChIP Low salt buffer (20 mM Tris/HCl pH 8.0, 2 mM EDTA, 150 mM NaCl, 1% (w/v) Triton X-100, 0.1% (w/v) SDS, 1 mM DTT), thrice with ChIP High salt buffer (20 mM Tris/HCl pH 8.0, 2 mM EDTA, 500 mM NaCl, 1% (w/v) Triton X-100, 0.1% (w/v) SDS, 1 mM DTT), once with LiCl buffer (10 mM Tris/HCl pH 8.0, 1 mM EDTA, 250 mM LiCl, 0.1% (w/v) NP-40, 1 mM DTT) and twice with TE buffer (10 mM Tris/HCl pH 8.0, 1 mM EDTA). Washing took place at 4 °C for 5 min with rotation followed by centrifugation (4 min, 400 g, 4 °C).

Precipitated material was eluted in 250 µl ChIP elution buffer (200 mM NaHCO3, 2% (w/v) SDS) for 45 min at RT with rotation followed by incubation at 95 °C for 15 min. The resin was precipitated by centrifugation for 3 min at 21,100 g and the supernatant was diluted 1:2 with nuclease free water. 14 µl of pre-cleared chromatin was added to 250 µl ChIP elution buffer and 250 µl nuclease free water as “input” sample. To separate protein-DNA crosslinks samples were treated with 40 µM NaCl over night at 65 °C with agitation. Upon addition of 40 mM Tris/HCl pH 6.8 and 1 mM EDTA samples were treated with 40 ng/μl Proteinase K (7528.1, Roth) for 1 h at 45 °C and DNA was purified using the QIAquick PCR purification kit (28106, Qiagen).

The following antibodies were used for precipitation: H3 (ab1791, Abcam), H4K16ac (07-329, Millipore), mouse IgG (12-371, Millipore).

Precipitated DNA was analyzed by high throughput sequencing.

### ChIP sequencing and data analysis

Precipitated DNA from six ChIP reactions was concentrated (Concentrator 5301, Eppendorf) and quantified using the Qubit dsDNA High-Sensitivity Assay Kit (Q32851, Thermo Fisher scientific). Sequencing libraries were generated from 0.5-2 ng of DNA using the MicroPlex Library Preparation Kit v2 (C05010012, Diagenode) according to manufacturer’s instructions. Quality of sequencing libraries was controlled on a Bioanalyzer 2100 using the Agilent High Sensitivity DNA Kit (Agilent). Pooled sequencing libraries were sequenced on the NextSeq 550 platform (Illumina) with 50 bases single reads.

Single-end reads were aligned to the *Drosophila melanogaster* genome (dm6) using Bowtie2 (2.5.3, Langmead et al., 2012) with default settings. Reads resulting from ChIP experiments using human spike-in chromatin were aligned to a composite genome (dm6 and hg38). Aligned reads were deduplicated using Samtools markdup (1.15.1, Danecek et al., 2021) and, in case of spike-in controlled experiments, separated according to genome origin using Split BAM (Samtools, 2.5.2). Scaling factors were calculated based on the number of human reads in ChIP from control and hangover knock-out cells. RPKM-normalized coverage vectors were generated using bamCoverage and bamCompare of Deeptools (3.5.4, Ramírez et al., 2016) with a bin size of 5. Here, reads were extended to an average fragment size of 500 bp (hangover & H4K16ac ChIP) or 200 bp (Rcd1 & MBD-R2 ChIP). Duplicates were ignored and spike-in scaling factors were included accordingly. Peaks were called against input control or H3 ChIP with MACS2 callpeak (2.2.9.1, Feng et al., 2012) and medium fragment size was estimated as 500 bp (hangover, H3 & H4K16ac ChIP) or 200 bp (Rcd1 & MBD-R2 ChIP). Peaks that lay less than 500 bp apart were fused using mergeBED.

To define common peaks between to BED files, peak sets were intersected with a default of 1 bp overlap. For hangover ‘high-confidence’ binding sites, intersecting peaks of replicate 1 with replicate 2 were calculated (-wa) and vice versa. Both intersections were combined and merged with a maximum distance of 500 bp on either strand. This way, for a given ‘high-confidence’ peak, the bigger region will be counted. Bedtools version 2.31.1 was used for all manipulations of BED files (Quinlan et al., 2010). Heatmaps were generated using Deeptools (3.5.4) and sorted according to average occupancy (defined as RPKM(ChIP)/RPKM(input)). Read counts were retrieved with MultiCovBed within Bedtools.

Metagene plots were generated over unique TSS retrieved from the UCSC table browser using bamCompare within Deeptools (ChIP over input, ratio of read number, bin size: 5). Peak distribution over genomic elements was determined using CEAS (1.0.0, Shin et al., 2009) within the Cistrome platform with default settings. To that end, hangover peaks were mapped to dm3 using the UCSC liftOver tool. Genomic elements were summarized as follows: promoter (promoter (<= 250 bp) + promoter (250-500 bp) + promoter (500-1000 bp)) and intergenic (distal intergenic + distal (<= 250 bp) + distal (250-500 bp) + distal (500-1000 bp)). Remaining definitions were adopted from CEAS.

Paired ATAC-seq reads were aligned as mentioned above. Mitochondrial reads, duplicates and pairs with a mapping quality < 30 were removed (Filter BAM 2.5.2; MarkDuplicates 3.1.1.0). Bedgraph files were generated using MACS2 (2.2.9.1; set extension size: 200; set shift size:-100) and converted to bigWig files using wigtobigwig (447, Kent et al., 2010).

Differential H4K16 acetylation was determined by comparing levels in hangover wild type (wt) and knock-out (ko) cells. Global differences in H4K16ac signal was evaluated by normalizing total H4K16ac and H3 read numbers to respective numbers of human spike-in reads. Acetylation was then defined as H4K16ac/H3 signal and normalized to hangover wt. TSS were recognized as H4K16 acetylated if a H4K16ac/H3 peak was found within a 1 kb window. H4K16ac and H3 reads in wt and hangover ko conditions were counted in these 1 kb intervals using MultiCovBed. H4K16ac counts were normalized to H3 counts and for each TSS a difference was calculated between wt and ko (ΔH4K16ac/H3). A frequency distribution of ΔH4K16ac/H3 was generated in Prism 10.0.3 with a bin width of 0.01. Mean (µ=0.00048) and standard deviation (#=0.12631) of ΔH4K16ac/H3 was determined and only those TSS were considered as differentially acetylated that lay outside of a µ±2# interval. Gene Ontology term analysis was performed using Metascape with “Express Analysis” settings (metascape.org; Zhou et at., 2019).

The following publicly available datasets were used: H3K4me3 (GSM3106567-GSM3106568, GSM3106577-GSM3106578, Tettey et al., 2019), H3K27ac and H3K27me3 (GSM2175510-GSM2175513, GSM2175500-GSM2175503, Rickels et al., 2016), ATAC-seq (GSM3381127, Albig et al., 2019), Rcd1 and MBD-R2 (E-MTAB-1085, Lam et al., 2012)

### RNA interference and RNA isolation

Sequences of double stranded RNA (dsRNA) and corresponding primers were designed using E-RNAi (<u>e-rnai.dkfz.de</u>; Horn et al., 2010) Double stranded RNA was generated using the MEGAscript T7 kit according to manufacturer’s instructions (AMB1334, Invitrogen). Templates for *in vitro* transcription were synthesized by PCR from S2[Cas9] cDNA using primers containing minimal T7 promoter sequences (Suppl. Table S1). For RNAi S2[Cas9] cells were seeded at 0.1-0.33 x10^6^ cells/ml and 5 µg/ml dsRNA was added. After four days cells were harvested for RNA or whole cell extract preparation.

RNA was isolated from S2[Cas9] cells using the peqGOLD Total RNA Kit (13-6834-02, VWR Life Science) with the peqGOLD DNase I Digest Kit (13-1091-01, VWR Life Science) according to manufacturer’s instructions.

### Protein extract preparation

For preparation of whole cell extracts cells were washed in PBS and subsequently lysed in RIPA buffer (50 mM Tris/HCl pH 8.0, 150 mM NaCl, 1 mM EDTA, 1 mM EGTA, 1% (w/v) NP-40, 0.5% (w/v) sodium deoxycholate, 0.1% (w/v) SDS, 10% (v/v) glycerol, 1 mM DTT). Suspension was incubated for 20 min at 4 °C with rotation followed by freeze/thaw lysis in liquid nitrogen. Cell debris were removed by centrifugation at 21,100 g at 4 °C for 30 min.

Nuclear proteins were isolated by a two-step lysis. After washing cells with PBS cells were lysed in hypotonic buffer B (10 mM Hepes/KOH pH 7.6, 10 mM KCl, 1.5 mM MgCl2, 1 mM DTT) for 20 min at 4 °C with rotation. Nuclei were isolated by centrifugation at 1,900 g for 20 min at 4 °C. The cytoplasmic fraction was discarded. Nuclei were resuspended in hypertonic buffer C (20 mM Hepes/KOH pH 7.6, 420 mM NaCl, 1.5 mM MgCl2, 0.2 mM EDTA, 20% (v/v) glycerol, 1 mM DTT) and proteins were extracted for 30 min at 4 °C with rotation. Lysate was cleared by centrifugation at 21,100 g for 30 min at 4 °C.

Protein concentrations were estimated against a BSA standard using a Bradford-based method (Protein Assay, 5000006, BioRad) according to manufacturer’s instructions.

Subcellular fractionation was performed using the Subcellular Protein Fractionation Kit for Cultured Cells (78840, Thermo Fisher Scientific) according to manufacturer’s instructions.

### Size exclusion chromatography

Nuclear extracts were diluted to a concentration of 4 mg/ml, 0.15 U/µl Benzonase (70664, Millipore) was added and the suspension was incubated at 4 °C overnight with rotation. Potential precipitates were removed by centrifugation at 21,000 g for 20 min. Extracts were loaded onto a Superose^TM^ 6 HR 10/30 GL column (17-0537-01, GE Healthcare) using an ÄKTA pure chromatography system (29018226, GE Healthcare) with a 200 µl injection loop. The column was operated in buffer EX300 (10 mM Hepes/KOH pH 7.6, 300 mM KCl, 1.5 mM MgCl_2_, 0.5 mM EGTA, 10% (v/v) glycerol, 1 mM DTT). 500 µl fractions were collected using a Fraction collector F9-R (29011362, GE Healthcare). Molecular weight across different fractions was calibrated using the Gel Filtration Cal Kit High Molecular Weight (28-4038-42, GE Healthcare).

To precipitate proteins in each fraction, 5 µl of StrataClean resin (400714, Agilent Technologies) was added and proteins were bound to the resin for 30 min at 4 °C with rotation. The resin was precipitated by centrifugation at 3,500 g for 5 min at 4 °C and proteins were eluted in 2x SDS loading buffer (100 mM Tris/HCl pH 6.8, 4% (w/v) SDS, 20% (v/v) glycerol, 0.2% (w/v) bromophenol blue, 200 mM DTT) for 5 min at 95 °C.

### Co-Immunoprecipitation of epitope-tagged proteins

Nuclear extracts were diluted to a final concentration of 1 mg/ml protein, 100 mM NaCl and 0.1% (w/v) NP-40 with buffers C-0 (20 mM Hepes/KOH pH 7.6, 1.5 mM MgCl_2_, 0.2 mM EDTA, 20% (v/v) glycerol, 0.131% (w/v) NP-40, 1 mM DTT) and C-100 (20 mM Hepes/KOH pH 7.6, 100 mM NaCl, 1.5 mM MgCl_2_, 0.2 mM EDTA, 20% (v/v) glycerol, 0.1% (w/v) NP-40, 1 mM DTT). 0.03 U/µl Benzonase (70664, Millipore) was added, diluted extracts were incubated at 4 °C for 1 h with rotation and contingent precipitates were removed by centrifugation (20 min, 5,000 g, 4 °C). GFP-Trap Agarose (gta, Proteintech) or ANTI-FLAG M2 Affinity Gel (A2220, Sigma) was washed in buffer C-100 and blocked with 1 mg/ml BSA and 1% (v/v) fish skin gelatin for 1 hour at 4 °C with rotation. 1-2 mg of diluted extract was added to 25 µl of resin and incubated overnight at 4 °C with rotation. After four washes with 1 ml IP150 buffer (25 mM Hepes/KOH pH 7.6, 150 mM NaCl, 12.5 mM MgCl_2_, 0.1 mM EDTA, 10% (v/v) glycerol 0.1% (w/v) NP-40, 1 mM DTT) proteins were eluted in SDS loading buffer (50 mM Tris/HCl pH 6.8, 2% (w/v) SDS, 10% (v/v) glycerol, 0.1% (w/v) bromophenol blue, 100 mM DTT) for 5 min at 95 °C. Purified proteins were analyzed by SDS-PAGE and Western Blot.

### Mass Spectrometry

Immunoprecipitation was performed from 10 mg of nuclear extract. Diluted extracts were added to 200 µl of non-blocked GFP-Trap Agarose (gta, Proteintech) or ANTI-FLAG M2 Affinity Gel (A2220, Sigma). Washing was performed on a Poly-Prep Chromatography Column (731-1550, BioRad) with 4x 10 ml IP150 buffer and twice with 100 mM NH_4_HCO_3_ buffer pH 7.5. Resin was removed from the column and bound proteins were analyzed by SDS-PAGE and silver staining or digested with trypsin for LC-MS analysis.

For protein quantification via LC-MS, beads were treated with 10 ng/l of trypsin in 1 M urea and 50 mM NH_4_HCO_3_ for 30 min, then rinsed with 50 mM NH_4_HCO_3_. Supernatants were digested overnight with 1 mM DTT. Before LC-MS analysis, peptides were alkylated and desalted. Peptides were then injected into a 25-cm analytical column (75 µm ID, 1.6 µm C18, Aurora-IonOpticks) with a 50-minute gradient from 2 to 35% acetonitrile in 0.1% formic acid. The effluent from the HPLC was electrosprayed directly into an Orbitrap Exploris 480 instrument operating in data-dependent mode to automatically transition between full-scan mass spectrometry (MS) and MS/MS acquisition. Typical mass spectrometric conditions were: spray voltage, 1.5 kV; no sheath and auxiliary gas flow; heated capillary temperature, 275 °C; ion selection threshold, 5×10^3^ counts; dynamic exclusion, 20s).

Specifically, Exploris 480 was set to: Survey full scan MS spectra (m/z 350–1400) were acquired with resolution 60000 at m/z 400 (max IT 40 ms, AGC target of 3×10^6^). The 15 most intense peptide ions with charge states between 2 and 6 were sequentially isolated to a target value of 2×10^5^, fragmented at 30% normalized collision energy and acquired with resolution 15000 at m/z 400 (max IT 40 ms).

MaxQuant 2.0.1.0 was used to identify and quantify proteins using iBAQ with the following parameters: DB: uniprot_AUP000000803_Dmelanogaster_20210325; MS tol: 10 ppm; MS/MS tol: 20 ppm Da; Peptide FDR: 0.1; Protein FDR: 0.01 min. peptide length: 7; Variable modifications: Oxidation (M); Fixed modifications: Carbamidomethyl (C); Peptides for protein quantitation: razor and unique; Min. peptides: 1; Min. ratio count: 2.

The quantified proteins (MaxQuant iBAQ Z-score normalised values) were compared using the adjusted t-test function from the Vulcano plot option from Perseus (missing values from the normal distribution replaced), width: 0.3 and downshift: 4, the false discovery rate (FDR): 0.05 and the S0 value: 0.1.

Proteins with a Δ log_2_ iBAQ (log_2_ iBAQ IP in control cells - log_2_ iBAQ IP in cells expressing GFP-or FLAG-tagged hangover) > 2 and a p-value < 0.05 were considered significantly enriched. Area-proportional Venn diagrams were generated using BioVenn (Hulsen et al., 2008). The STRING database (string-db.org, version 12.0) was used to generate a physical subnetwork based on experiments and databases. Proteins were grouped using MCL clustering (inflation parameter: 3). Gene Ontology term analysis was performed using Metascape with “Express Analysis” settings (metascape.org; Zhou et at., 2019).

### SDS-PAGE, Western Blot and Silver staining

SDS-polyacrylamide gel electrophoreses was used to separate proteins for subsequent Western blotting. Here, proteins were transferred to polyvinylidene difluride (PVDF) membranes (T830.1, Roth) in Pierce Western Blot Transfer Buffer (35040, Thermo Fisher Scientific). Blocking was conducted for 1 hour at room temperature in Blocking buffer (PBS, 0.1% (w/v) Tween-20, 5% (w/v) non-fat dry milk) and primary antibodies were applied in Blocking buffer over night at 4 °C. Membranes were washed four times in washing buffer (PBS, 0.1% Tween-20) and HRP-coupled secondary antibodies were applied in Blocking buffer for 2 h at room temperature (anti-mouse IgG (NA931, GE Healthcare), anti-rabbit IgG (NA934, GE Healthcare), anti-rat IgG (31470, Thermo Fisher Scientific)). Again, membranes were washed four times in washing buffer and chemiluminescence was detected using Immobilon Western Blot Chemiluminescence HRP substrate (WBKLS0500, Millipore) on a ChemiDoc^TM^ Touch Imaging System from Bio-Rad.

The following antibodies and antisera were used for detection: hangover (1:5,000; Scholz et al., 2005), GFP (1:5,000; clone [3H9] from Chromotek), FLAG (1:8,000; clone M2 from Sigma), FLAG (1:8,000; F7425 from Sigma), Tubulin beta (1:8,000; clone KMX-1 from Merck Millipore), Lamin Dm0 (1:5,000; clone ADL67.10 from DHSB), H3 (1:10,000; ab1791 from Abcam), nsl1 (1:5,000; Mendjan et al., 2006), MBD-R2 (1:5,000; Mendjan et al., 2006), mof (1:12,000; Mendjan et al., 2006), mle (1:12,000; Mendjan et al., 2006), Rcd5 (1:5,000; Gaub et al., 2020).

For silver staining of purified proteins, samples were supplemented with NuPAGE^TM^ LDS sample buffer (NP0007, Invitrogen), loaded on NuPAGE^TM^ 4-12% Bis-Tris gradient gels (NP0321, Invitrogen) and electrophoresed in NuPAGE^TM^ MOPS SDS running buffer (NP0001, Invitrogen). Gradient gels were then stained with the SilverQuest Silver Staining Kit (LC6070, Invitrogen) according to manufacturer’s instructions.

### qPCR

1 µg of total RNA was utilized for cDNA preparation using the SensiFAST cDNA Synthesis Kit (BIO-65054, Bioline) and the reaction was subsequently diluted 1:10. Eluted DNA from ChIP experiments was diluted 1:4. Diluted DNA samples were measured by qPCR using the SensiFast SYBR Lo-ROX Kit (BIO-94050, Bioline) according to manufacturer’s instructions. Each sample was measured in triplicates on a Stratagene Mx3000P thermocycler (Agilent Technologies). Sequences of primers used for amplification can be found in table S1.

For expression analysis using RT-qPCR, samples were first normalized to the control condition (hangover wt with dsLuc) and then scaled according to a normalization factor composed of the geometric mean of three reference genes (Act5C, βTub56D and RpL32). For that, the following calculations were applied:

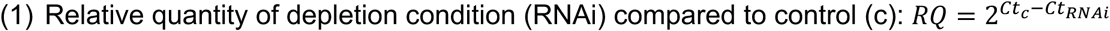

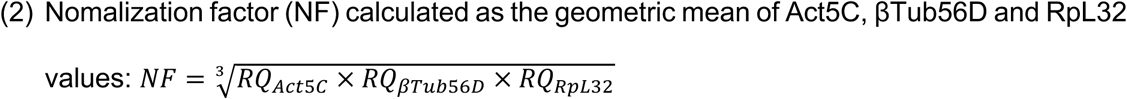

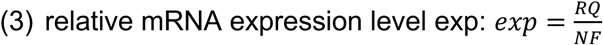

Error bars represent the standard deviation of five independent transfection experiments.

### RNA sequencing and data analysis

RNA quality was assessed using the Experion RNA StdSens Analysis Kit (BioRad). RNASeq libraries were prepared from total RNA using the TruSeq Stranded mRNA kit (Illumina) according to manufacturer’s instructions. Quality of sequencing libraries was controlled on a Bioanalyzer 2100 using the Agilent High Sensitivity DNA Kit (Agilent). Pooled sequencing libraries were quantified and sequenced on the NextSeq 550 platform (Illumina) with 50 bases single reads.

Read quality was assessed with FastQC (0.74) and adaptor sequences and low-quality bases were removed using Trimmomatic (0.39, Bolger et al., 2014). Single-end or paired-end reads were mapped to the *Drosophila melanogaster* genome (dm6) using RNA Star with 2-pass mapping (2.7.11a, Dobin et al., 2013). Reads were counted with featureCounts (default settings, 2.0.3, Liao et al., 2014), using the dmel_r6.56 GTF annotation file from flybase.org.

Fold change values and normalized counts were determined using DESeq2 (2.11.40.8, Love et al., 2014). A gene was defined as “regulated” when its adjusted p-value was lower than 0.05.

To intersect regulated genes with genomic binding sites, gene locations were retrieved from flybase.org (FB2024_2). To include promoter regions, locations were expanded by adding 500 bp upstream of the TSS. These genomic regions were intersected with hangover binding sites using bedtools (default settings, 2.31.1, Quinlan et al., 2010). Heatmaps were generated on fold change values that were scaled by row using heatmap2 (3.1.3.1). Gene set enrichment analysis was performed using GSEA (4.3.3, Subramanian, Tamayo et al., 2005, Mootha, Lindgren et al., 2003) with custom gene sets.

The following publicly available datasets were used: NSL1 RNAi (GSM3344618-GSM3344623, Lam et al., 2019), NSL1 & NSL3 RNAi (GSM4029652-GSM4029654, GSM4029657-GSM4029659, GSM4029662-GSM4029664, Gaub et al., 2020).

### Software, analysis platforms and data availability

The following software was used for analysis and representation of data: Image Lab (version 6.1.0 build 7, Bio-Rad), GraphPad Prism (version 10.0.3), Perseus (version 1.6.15.0, Tayanova, Temu et al., 2016). Genome-wide data was analyzed on the following Galaxy web platforms (Abueg et al., 2024): Galaxy Europe (usegalaxy.eu), Galaxy (usegalaxy.org), Galaxy/Cistrome (cistrome.org/ap/root).

Data generated in this study is available from the following repositories: European Nucleotide Archive (ebi.ac.uk/ena/browser/home, accession number: PRJEB88170) & ProteomeXchange Consortium via the PRIDE partner repository (Perez-Riverol et al., 2025; Deutsch et al., 2023; https://www.ebi.ac.uk/pride/, accession number: PXD062946).

## Acknowledgements

We thank Asifa Akhtar, Henrike Scholz and Klaus Förstemann for the generous gift of antibodies, plasmids and cell lines and Sandra B. Hake for critically reading the manuscript.

## Funding

JL and AB were funded by the Deutsche Forschungsgemeinschaft (DFG, German Research Foundation) - TRR81 A01 & BR2102/8.

## Author Contributions

JL devised and carried out experiments, analyzed results, performed bioinformatic analyses and wrote the manuscript, LS carried out and analyzed experiments, IF and AI analyzed protein samples with mass spectrometry and analyzed results, AN and TS performed high throughput sequencing, AB devised experiments, analyzed results and wrote the manuscript.

---

**Supplementary Figure 1.**
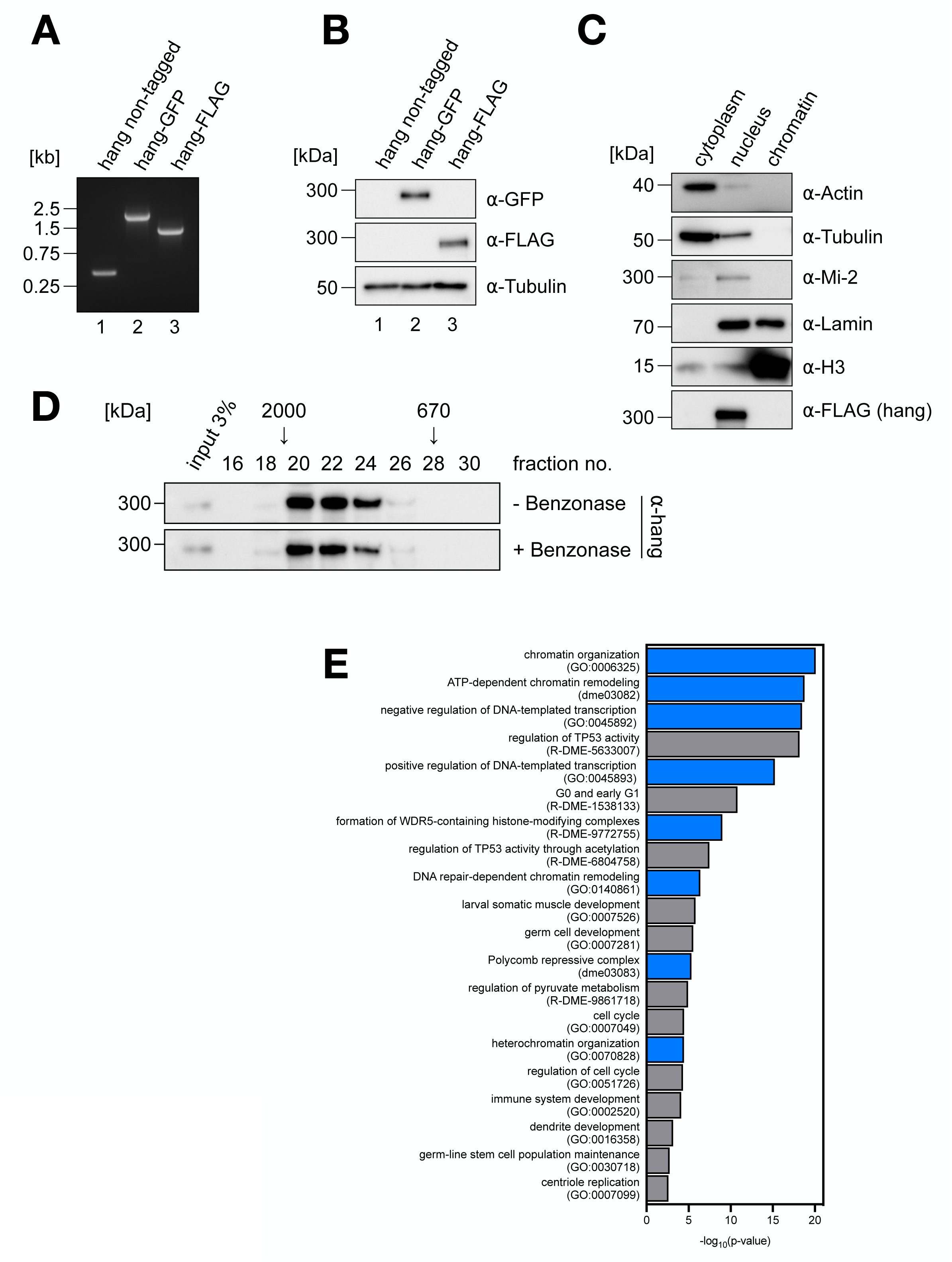
A. PCR amplification of genomic DNA from control S2 cells (hang non-tagged, lane 1) and cells modified to express C-terminally epitope-tagged hangover (hang-GFP, lane 2; hang-FLAG, lane 3). PCR was performed using primers flanking insertion site at the 3’ end of the hangover open reading frame resulting in 383 bp for the non-tagged, 1921 bp for a GFP-tagged and 1261 bp for a FLAG-tagged allele. DNA size markers are indicated on the left. **B** Western blot of nuclear extracts from control S2 cells (hang non-tagged, lane 1) and cells expressing hang-GFP (lane 2) or hang-FLAG (lane 3). Expression of tagged proteins was verified using GFP and FLAG antibodies. Tubulin served as loading control. Molecular weight markers are indicated on the left. **C** Western blot of subcellular fractions from hang-FLAG expressing S2 cells. Purity of fractions was monitored with antibodies against Actin & Tubulin (cytoplasm), Mi-2 (nucleoplasm) and H3 (chromatin). Lamin is found in both the nuclear and the chromatin fraction. Hang-FLAG was detected with a FLAG antibody. Molecular weight marker bands are indicated on the left. **D** Size exclusion chromatography profile of hangover. Prior to chromatography, nuclear extracts from S2 cells were incubated with or without Benzonase overnight. Void volume was estimated at fraction 19. Molecular weight markers on top indicate the approximate elution size. **E** Gene Ontology term analysis of hangover high-confidence interactors. GO terms associated with chromatin and transcription are highlighted in blue.

**Supplementary Figure 3.**
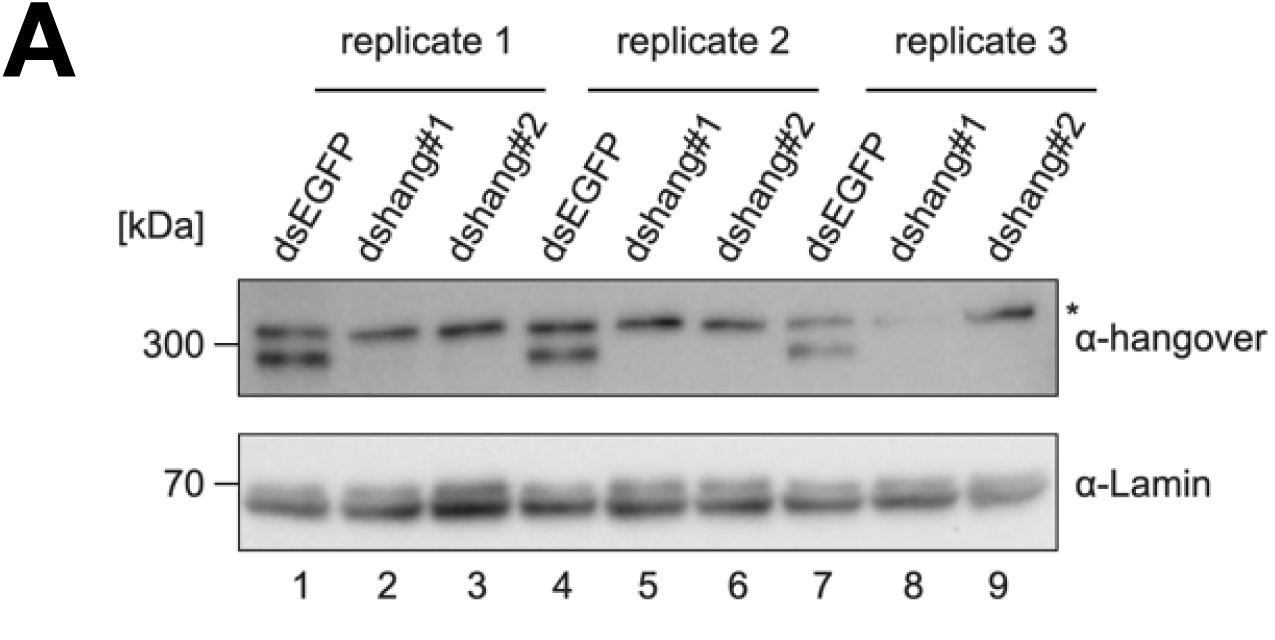
A. Western blot of S2 whole cell extracts upon treatment with control dsRNA (dsEGFP) and dsRNA targeting hangover (dshang#1 & dshang#2). Extracts were prepared from three independent transfections that were used for RNA-seq (replicates 1, 2 and 3). Membranes were probed with anti-hangover and anti-Lamin antibodies (loading control). Asterisk marks a non-specific band. Molecular weight markers are indicated on the left.

**Supplementary Figure 4.**
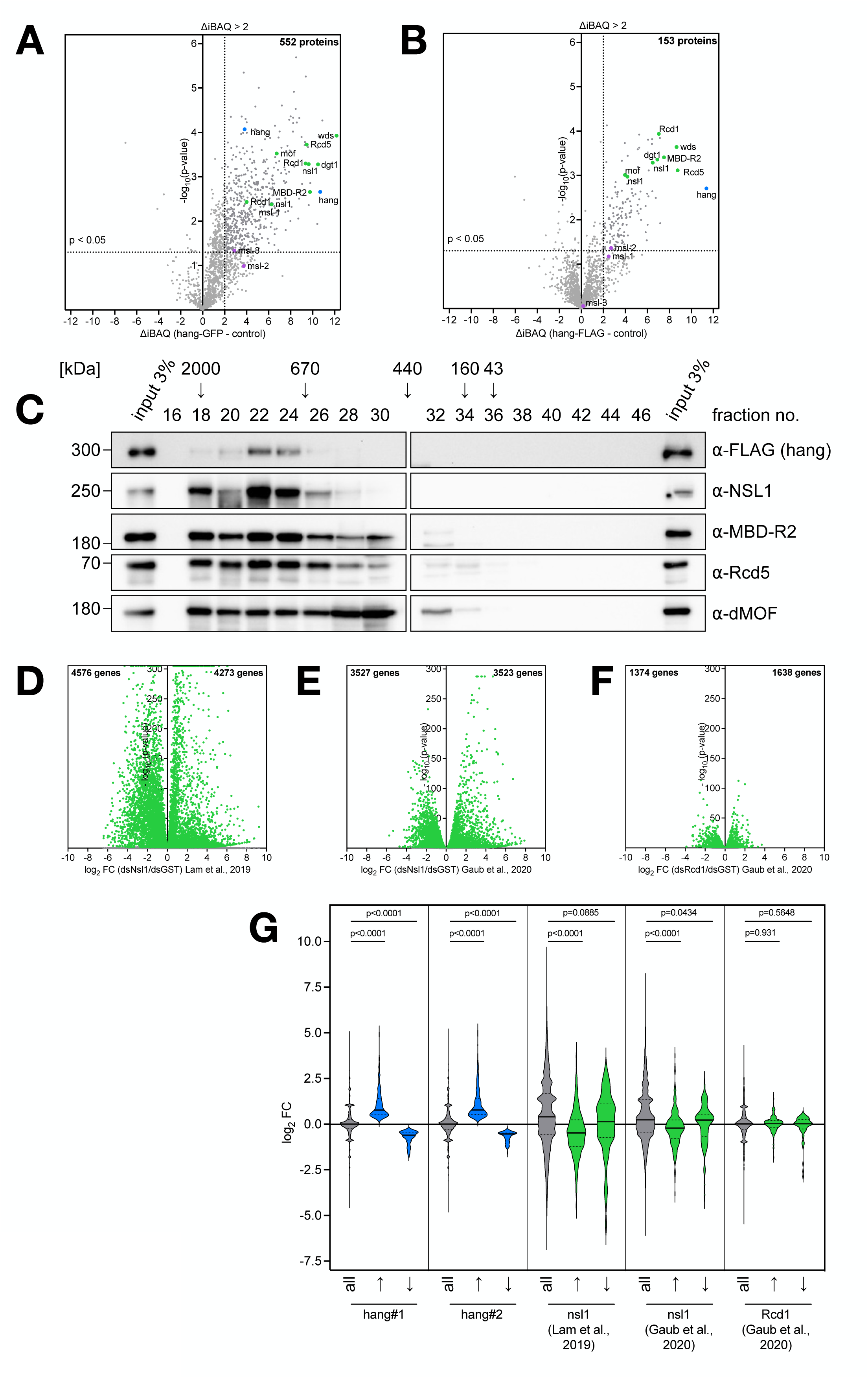
A &. **B** ⍺-GFP-(**A**) and ⍺-FLAG IP (**B**) IP-MS/MS results represented as volcano plots. Proteins that surpassed Δ log_2_ iBAQ and p-value cutoffs (Δ log_2_ iBAQ>2, p<0.05; represented by dotted lines) are considered significantly enriched. The number of these proteins is indicated above. Hangover is marked in blue, NSL complex subunits in green and MSL subunits in purple. **C** Size exclusion chromatography of nuclear extracts of S2 cells expressing FLAG-tagged hangover. Fractions were probed with antibodies against FLAG and NSL complex subunits (depicted on the right). Void volume was estimated at fraction 18. Molecular weight markers on top indicate the approximate elution size. **D-F** Volcano plots of genes deregulated upon RNAi-mediated depletion of NSL complex subunits. Log_2_ fold changes of are shown for nsl1 dsRNA treated vs. dsGST treated cells (**D & E**, Lam et al., 2019 and Gaub et al., 2020, respectively) and dsRcd1 vs. dsGST treated cells (**F**, Gaub et al., 2020). Genes with p<0.05 are marked in green and the number of these genes is indicated above. **G** Violin plots depicting the distribution of log_2_ fold changes upon depletion of hangover (sections 1 & 2) and NSL complex subunits (sections 3-5). Log_2_ FC were evaluated on all genes (all) and genes with increased (↑) or decreased (↓) expression upon hangover depletion. Solid lines represent median FC and dotted lines represent quartiles. Distributions were compared using two-tailed Mann-Whitney statistics and p-values are indicated on top.

**Supplementary Figure 5.**
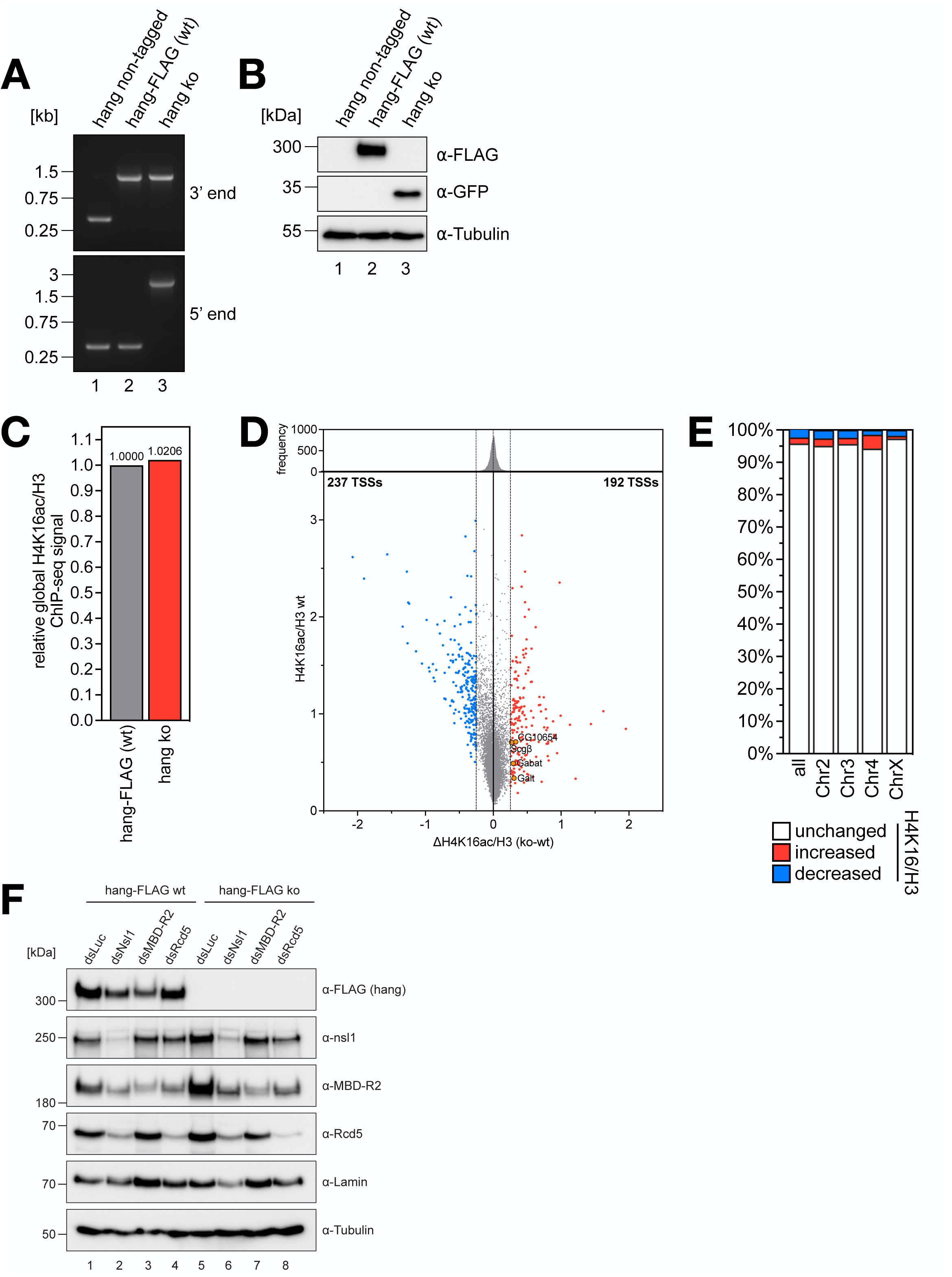
A. PCR amplification of genomic DNA from control S2 cells (hang non-tagged, lane 1), cells modified to express C-terminally FLAG-tagged hangover (lane 2) or hangover knock-out cells (lane 3). PCR was performed using primers at the 3’ end of the hangover open reading frame to test integration of the FLAG-tag sequence resulting in 383 bp for the non-tagged and 1261 bp for a FLAG-tagged allele. Primers at the 5’ end were used to verify integration of the GFP expression cassette resulting in 395 bp for a wt and 2151 bp for a ko allele. DNA size markers are indicated on the left. **B** Western blot of nuclear extracts from control S2 cells (hang non-tagged, lane 1) and cells expressing hang-FLAG (lane 2) or hangover knock-out cells (lane 3). Expression of tagged hangover was verified using a FLAG antibody. Successful integration of the GFP expression cassette and therefore disruption of the hangover locus was tested using a GFP antibody. Tubulin served as loading control. Molecular weight markers are indicated on the left. **C** Quantification of global H4K16ac ChIP-seq signal in hangover wt and ko cells. Total mapped H4K16ac and H3 reads were normalized to their respective human spike-in read number. H4K16ac signal is expressed as H4K16ac/H3 relative to signal in hangover wt cells. **D** Differential H4K16ac at TSS upon hangover ko. H4K16ac/H3 signal in wt cells is plotted against the difference of H4K16ac/H3 between wt and hangover ko cells (ΔH4K16ac/H3). Frequency distribution of ΔH4K16ac/H3 is shown above. TSS with increased H4K16ac are highlighted in red, TSS with decreased H4K16ac are highlighted in blue and their numbers are indicated above. Dotted lines represent the µ±2# interval. TSS of CG10654, Scgβ, Galt and Gabat are marked in orange. **E** Distribution of differentially H4K16 acetylated TSS over different chromosomes, expressed as percent of all analyzed TSS on the respective chromosome. Fractions of TSS with increased H4K16ac are marked in red. Fractions of TSS with decreased H4K16ac are marked in blue. **F** Representative Western blot of nuclear extracts from wt and hangover ko cells treated with control dsRNA (dsLuc) or dsRNA against nsl1, MBD-R2 and Rcd5. Hang-FLAG expression was confirmed using a FLAG antibody and RNAi efficiency was monitored using antibodies indicated on the right. Lamin and Tubulin served as loading controls. Molecular weight markers are indicated on the left.

## Notes

### Competing Interest Statement

The authors have declared no competing interest.

## References

Teves, S. S., & Henikoff, S. (2011). Heat shock reduces stalled RNA polymerase II and nucleosome turnover genome-wide. Genes and Development, 25(22), 2387–2397. 10.1101/gad.177675.111

Guertin, M. J., & Lis, J. T. (2010). Chromatin landscape dictates HSF binding to target DNA elements. PLoS Genetics, 6(9). 10.1371/journal.pgen.1001114

Teves, S. S., & Henikoff, S. (2013). The heat shock response: A case study of chromatin dynamics in gene regulation. In Biochemistry and Cell Biology (Vol. 91, Issue 1, pp. 42–48). 10.1139/bcb-2012-0075

Panniers, R. (1994). Translational control during heat shock. Biochimie, 76(8), 737–747. 10.1016/0300-9084(94)90078-7

Elvira, R., Cha, S. J., Noh, G. M., Kim, K., & Han, J. (2020). PERK-mediated eIF2α phosphorylation contributes to the protection of dopaminergic neurons from chronic heat stress in Drosophila. International Journal of Molecular Sciences, 21(3). 10.3390/ijms21030845

Saxton, R. A., & Sabatini, D. M. (2017). mTOR Signaling in Growth, Metabolism, and Disease. In Cell (Vol. 168, Issue 6, pp. 960–976). Cell Press. 10.1016/j.cell.2017.02.004

Scholz, H., Franz, M., & Heberlein, U. (2005). The hangover gene defines a stress pathway required for ethanol tolerance development. Nature, 436(7052), 845–847. 10.1038/nature03864

Schwenkert, I., Eltrop, R., Funk, N., Steinert, J. R., Schuster, C. M., & Scholz, H. (2008). The hangover gene negatively regulates bouton addition at the Drosophila neuromuscular junction. Mechanisms of Development, 125(8), 700–711. 10.1016/j.mod.2008.04.004

Ruppert, M., Franz, M., Saratsis, A., Velo Escarcena, L., Hendrich, O., Gooi, L. M., Schwenkert, I., Klebes, A., & Scholz, H. (2017). Hangover Links Nuclear RNA Signaling to cAMP Regulation via the Phosphodiesterase 4d Ortholog dunce. Cell Reports, 18(2), 533–544. 10.1016/j.celrep.2016.12.048

Cassandri, M., Smirnov, A., Novelli, F., Pitolli, C., Agostini, M., Malewicz, M., Melino, G., & Raschellà, G. (2017). Zinc-finger proteins in health and disease. In Cell Death Discovery (Vol. 3, Issue 1). Springer Nature. 10.1038/cddiscovery.2017.71

Brayer, K. J., & Segal, D. J. (2008). Keep your fingers off my DNA: Protein-protein interactions mediated by C2H2 zinc finger domains. In Cell Biochemistry and Biophysics (Vol. 50, Issue 3, pp. 111–131). 10.1007/s12013-008-9008-5

Sheikh, B. N., Guhathakurta, S., & Akhtar, A. (2019). The non-specific lethal (NSL) complex at the crossroads of transcriptional control and cellular homeostasis. EMBO Reports, 20(7). 10.15252/embr.201847630

Gelbart, M. E., Larschan, E., Peng, S., Park, P. J., & Kuroda, M. I. (2009). Drosophila MSL complex globally acetylates H4K16 on the male X chromosome for dosage compensation. Nature Structural and Molecular Biology, 16(8), 825–832. 10.1038/nsmb.1644

Lam, K. C., Mühlpfordt, F., Vaquerizas, J. M., Raja, S. J., Holz, H., Luscombe, N. M., Manke, T., & Akhtar, A. (2012). The NSL complex regulates housekeeping genes in Drosophila. PLoS Genetics, 8(6). 10.1371/journal.pgen.1002736

Lam, K. C., Chung, H. R., Semplicio, G., Iyer, S. S., Gaub, A., Bhardwaj, V., Holz, H., Georgiev, P., & Akhtar, A. (2019). The NSL complex-mediated nucleosome landscape is required to maintain transcription fidelity and suppression of transcription noise. Genes and Development, 33(7–8), 452–465. 10.1101/gad.321489.118

Gaub, A., Sheikh, B. N., Basilicata, M. F., Vincent, M., Nizon, M., Colson, C., Bird, M. J., Bradner, J. E., Thevenon, J., Boutros, M., & Akhtar, A. (2020). Evolutionary conserved NSL complex/BRD4 axis controls transcription activation via histone acetylation. Nature Communications, 11(1). 10.1038/s41467-020-16103-0

Leader, D. P., Krause, S. A., Pandit, A., Davies, S. A., & Dow, J. A. T. (2018). FlyAtlas 2: A new version of the Drosophila melanogaster expression atlas with RNA-Seq, miRNA-Seq and sex-specific data. Nucleic Acids Research, 46(D1), D809–D815. 10.1093/nar/gkx976

Krause, S. A., Overend, G., Dow, J. A. T., & Leader, D. P. (2022). FlyAtlas 2 in 2022: Enhancements to the Drosophila melanogaster expression atlas. Nucleic Acids Research, 50(D1), D1010–D1015. 10.1093/nar/gkab971

Oksuz, O., Henninger, J. E., Warneford-Thomson, R., Zheng, M. M., Erb, H., Vancura, A., Overholt, K. J., Hawken, S. W., Banani, S. F., Lauman, R., Reich, L. N., Robertson, A. L., Hannett, N. M., Lee, T. I., Zon, L. I., Bonasio, R., & Young, R. A. (2023). Transcription factors interact with RNA to regulate genes. Molecular Cell, 83(14), 2449–2463.e13. 10.1016/j.molcel.2023.06.012

Erokhin, M. M., Shidlovskii, Y. v., Lomaev, D. v., Georgiev, P. G., & Chetverina, D. A. (2021). Sfmbt Co-purifies with Hangover and SWI/SNF-Remodelers in Drosophila melanogaster. Doklady Biochemistry and Biophysics, 500(1), 304–307. 10.1134/S1607672921050069

Kunert, N., Wagner, E., Murawska, M., Klinker, H., Kremmer, E., & Brehm, A. (2009). dMec: A novel Mi-2 chromatin remodelling complex involved in transcriptional repression. EMBO Journal, 28(5), 533–544. 10.1038/emboj.2009.3

Clapier, C. R., & Cairns, B. R. (2009). The biology of chromatin remodeling complexes. In Annual Review of Biochemistry (Vol. 78, pp. 273–304). 10.1146/annurev.biochem.77.062706.153223

Spain, M. M., Caruso, J. A., Swaminathan, A., & Pile, L. A. (2010). Drosophila SIN3 isoforms interact with distinct proteins and have unique biological functions. Journal of Biological Chemistry, 285(35), 27457–27467. 10.1074/jbc.M110.130245

Partridge, E. C., Chhetri, S. B., Prokop, J. W., Ramaker, R. C., Jansen, C. S., Goh, S. T., Mackiewicz, M., Newberry, K. M., Brandsmeier, L. A., Meadows, S. K., Messer, C. L., Hardigan, A. A., Coppola, C. J., Dean, E. C., Jiang, S., Savic, D., Mortazavi, A., Wold, B. J., Myers, R. M., & Mendenhall, E. M. (2020). Occupancy maps of 208 chromatin-associated proteins in one human cell type. Nature, 583(7818), 720–728. 10.1038/s41586-020-2023-4

Malovannaya, A., Lanz, R. B., Jung, S. Y., Bulynko, Y., Le, N. T., Chan, D. W., Ding, C., Shi, Y., Yucer, N., Krenciute, G., Kim, B. J., Li, C., Chen, R., Li, W., Wang, Y., O’Malley, B. W., & Qin, J. (2011). Analysis of the human endogenous coregulator complexome. Cell, 145(5), 787–799. 10.1016/j.cell.2011.05.006

Kahn, T. G., Savitsky, M., Kuong, C., Jacquier, C., Cavalli, G., Chang, J.-M., & Schwartz, Y. B. (2023). Topological screen identifies hundreds of Cp190-and CTCF-dependent Drosophila chromatin insulator elements. https://www.science.org

Bhattacharya, M., Lyda, S. F., & Lei, E. P. (2024). Chromatin insulator mechanisms ensure accurate gene expression by controlling overall 3D genome organization. In Current Opinion in Genetics and Development (Vol. 87). Elsevier Ltd. 10.1016/j.gde.2024.102208

Sheikh, B. N., Bechtel-Walz, W., Lucci, J., Karpiuk, O., Hild, I., Hartleben, B., Vornweg, J., Helmstädter, M., Sahyoun, A. H., Bhardwaj, V., Stehle, T., Diehl, S., Kretz, O., Voss, A. K., Thomas, T., Manke, T., Huber, T. B., & Akhtar, A. (2016). MOF maintains transcriptional programs regulating cellular stress response. Oncogene, 35(21), 2698–2710. 10.1038/onc.2015.335

Schmitges, F. W., Radovani, E., Najafabadi, H. S., Barazandeh, M., Campitelli, L. F., Yin, Y., Jolma, A., Zhong, G., Guo, H., Kanagalingam, T., Dai, W. F., Taipale, J., Emili, A., Greenblatt, J. F., & Hughes, T. R. (2016). Multiparameter functional diversity of human C2H2 zinc finger proteins. Genome Research, 26(12), 1742–1752. 10.1101/gr.209643.116

## References (Methods)

Böttcher, R., Hollmann, M., Merk, K., Nitschko, V., Obermaier, C., Philippou-Massier, J., Wieland, I., Gaul, U., & Förstemann, K. (2014). Efficient chromosomal gene modification with CRISPR/cas9 and PCR-based homologous recombination donors in cultured Drosophila cells. Nucleic Acids Research, 42(11). 10.1093/nar/gku289

Lenz, J., Liefke, R., Funk, J., Shoup, S., Nist, A., Stiewe, T., Schulz, R., Tokusumi, Y., Albert, L., Raifer, H., Förstemann, K., Vázquez, O., Tokusumi, T., Fossett, N., & Brehm, A. (2021). Ush regulates hemocyte-specific gene expression, fatty acid metabolism and cell cycle progression and cooperates with dNuRD to orchestrate hematopoiesis. In PLoS Genetics (Vol. 17, Issue 2). 10.1371/JOURNAL.PGEN.1009318

Langmead, B., & Salzberg, S. L. (2012). Fast gapped-read alignment with Bowtie 2. Nature Methods, 9(4), 357–359. 10.1038/nmeth.1923

Danecek, P., Bonfield, J. K., Liddle, J., Marshall, J., Ohan, V., Pollard, M. O., Whitwham, A., Keane, T., McCarthy, S. A., & Davies, R. M. (2021). Twelve years of SAMtools and BCFtools. GigaScience, 10(2). 10.1093/gigascience/giab008

Ramírez, F., Ryan, D. P., Grüning, B., Bhardwaj, V., Kilpert, F., Richter, A. S., Heyne, S., Dündar, F., & Manke, T. (2016). deepTools2: a next generation web server for deep-sequencing data analysis. Nucleic Acids Research, 44(W1), W160–W165. 10.1093/NAR/GKW257

Feng, J., Liu, T., Qin, B., Zhang, Y., & Liu, X. S. (2012). Identifying ChIP-seq enrichment using MACS. Nature Protocols, 7(9), 1728–1740. 10.1038/nprot.2012.101

Quinlan, A. R., & Hall, I. M. (2010). BEDTools: A flexible suite of utilities for comparing genomic features. Bioinformatics, 26(6), 841–842. 10.1093/bioinformatics/btq033

Shin, H., Liu, T., Manrai, A. K., & Liu, S. X. (2009). CEAS: Cis-regulatory element annotation system. Bioinformatics, 25(19), 2605–2606. 10.1093/bioinformatics/btp479

Kent, W. J., Zweig, A. S., Barber, G., Hinrichs, A. S., & Karolchik, D. (2010). BigWig and BigBed: Enabling browsing of large distributed datasets. Bioinformatics, 26(17), 2204–2207. 10.1093/bioinformatics/btq351

Zhou, Y., Zhou, B., Pache, L., Chang, M., Khodabakhshi, A. H., Tanaseichuk, O., Benner, C., & Chanda, S. K. (2019). Metascape provides a biologist-oriented resource for the analysis of systems-level datasets. Nature Communications, 10(1). 10.1038/s41467-019-09234-6

Tettey, T. T., Gao, X., Shao, W., Li, H., Story, B. A., Chitsazan, A. D., Glaser, R. L., Goode, Z. H., Seidel, C. W., Conaway, R. C., Zeitlinger, J., Blanchette, M., & Conaway, J. W. (2019). A Role for FACT in RNA Polymerase II Promoter-Proximal Pausing. Cell Reports, 27(13), 3770–3779.e7. 10.1016/j.celrep.2019.05.099

Rickels, R., Hu, D., Collings, C. K., Woodfin, A. R., Piunti, A., Mohan, M., Herz, H. M., Kvon, E., & Shilatifard, A. (2016). An Evolutionary Conserved Epigenetic Mark of Polycomb Response Elements Implemented by Trx/MLL/COMPASS. Molecular Cell, 63(2), 318–328. 10.1016/j.molcel.2016.06.018

Albig, C., Tikhonova, E., Krause, S., Maksimenko, O., Regnard, C., & Becker, P. B. (2019). Factor cooperation for chromosome discrimination in Drosophila. Nucleic Acids Research, 47(4), 1706–1724. 10.1093/nar/gky1238

Horn, T., & Boutros, M. (2010). E-RNAi: A web application for the multi-species design of RNAi reagents-2010 update. Nucleic Acids Research, 38(SUPPL. 2). 10.1093/nar/gkq317

Hulsen, T., de Vlieg, J., & Alkema, W. (2008). BioVenn - A web application for the comparison and visualization of biological lists using area-proportional Venn diagrams. BMC Genomics, 9. 10.1186/1471-2164-9-488

Mendjan, S., Taipale, M., Kind, J., Holz, H., Gebhardt, P., Schelder, M., Vermeulen, M., Buscaino, A., Duncan, K., Mueller, J., Wilm, M., Stunnenberg, H. G., Saumweber, H., & Akhtar, A. (2006). Nuclear pore components are involved in the transcriptional regulation of dosage compensation in Drosophila. Molecular Cell, 21(6), 811–823. 10.1016/j.molcel.2006.02.007

Bolger, A. M., Lohse, M., & Usadel, B. (2014). Trimmomatic: A flexible trimmer for Illumina sequence data. Bioinformatics, 30(15), 2114–2120. 10.1093/bioinformatics/btu170

Dobin, A., Davis, C. A., Schlesinger, F., Drenkow, J., Zaleski, C., Jha, S., Batut, P., Chaisson, M., & Gingeras, T. R. (2013). STAR: Ultrafast universal RNA-seq aligner. Bioinformatics, 29(1), 15–21. 10.1093/bioinformatics/bts635

Liao, Y., Smyth, G. K., & Shi, W. (2014). FeatureCounts: An efficient general purpose program for assigning sequence reads to genomic features. Bioinformatics, 30(7), 923–930. 10.1093/bioinformatics/btt656

Love, M. I., Huber, W., & Anders, S. (2014). Moderated estimation of fold change and dispersion for RNA-seq data with DESeq2. Genome Biology, 15(12). 10.1186/s13059-014-0550-8

Subramanian, A., Tamayo, P., Mootha, V. K., Mukherjee, S., Ebert, B. L., Gillette, M. A., Paulovich, A., Pomeroy, S. L., Golub, T. R., Lander, E. S., & Mesirov, J. P. (2005). Gene set enrichment analysis: A knowledge-based approach for interpreting genome-wide expression profiles. Proc Natl Acad Sci U S A, 102(43), 15545–15550. www.pnas.orgcgidoi10.1073pnas.0506580102

Mootha, V. K., Lindgren, C. M., Eriksson, K.-F., Subramanian, A., Sihag, S., Lehar, J., Puigserver, P., Carlsson, E., Ridderstråle, M., Laurila, E., Houstis, N., Daly, M. J., Patterson, N., Mesirov, J. P., Golub, T. R., Tamayo, P., Spiegelman, B., Lander, E. S., Hirschhorn, J. N.,… Groop, L. C. (2003). PGC-1α-responsive genes involved in oxidative phosphorylation are coordinately downregulated in human diabetes. In NATURE GENETICS VOLUME (Vol. 34, Issue 3). http://www.nature.com/naturegenetics

Tyanova, S., Temu, T., Sinitcyn, P., Carlson, A., Hein, M. Y., Geiger, T., Mann, M., & Cox, J. (2016). The Perseus computational platform for comprehensive analysis of (prote)omics data. In Nature Methods (Vol. 13, Issue 9, pp. 731–740). Nature Publishing Group. 10.1038/nmeth.3901

Abueg, L. A. L., Afgan, E., Allart, O., Awan, A. H., Bacon, W. A., Baker, D., Bassetti, M., Batut, B., Bernt, M., Blankenberg, D., Bombarely, A., Bretaudeau, A., Bromhead, C. J., Burke, M. L., Capon, P. K., Čech, M., Chavero-Díez, M., Chilton, J. M., Collins, T. J.,… Zoabi, R. (2024). The Galaxy platform for accessible, reproducible, and collaborative data analyses: 2024 update. Nucleic Acids Research, 52(W1), W83–W94. 10.1093/nar/gkae410

Perez-Riverol, Y., Bandla, C., Kundu, D. J., Kamatchinathan, S., Bai, J., Hewapathirana, S., John, N. S., Prakash, A., Walzer, M., Wang, S., & Vizcaíno, J. A. (2025). The PRIDE database at 20 years: 2025 update. Nucleic Acids Research, 53(D1), D543–D553. 10.1093/nar/gkae1011

Deutsch, E. W., Bandeira, N., Perez-Riverol, Y., Sharma, V., Carver, J. J., Mendoza, L., Kundu, D. J., Wang, S., Bandla, C., Kamatchinathan, S., Hewapathirana, S., Pullman, B. S., Wertz, J., Sun, Z., Kawano, S., Okuda, S., Watanabe, Y., Maclean, B., Maccoss, M. J.,… Vizcaíno, J. A. (2023). The ProteomeXchange consortium at 10 years: 2023 update. Nucleic Acids Research, 51(D1), D1539–D1548. 10.1093/nar/gkac1040

